# An Enhancer Trap system to track developmental dynamics in Marchantia polymorpha

**DOI:** 10.1101/2023.07.06.547788

**Authors:** Alan O. Marron, Susana Sauret-Gueto, Marius Rebmann, Linda Silvestri, Marta Tomaselli, Jim Haseloff

**Affiliations:** Department of Plant Sciences, University of Cambridge, Downing Street, Cambridge CB2 3EA, UK; Crop Science Centre, University of Cambridge, 93 Lawrence Weaver Road, Cambridge CB3 0LE, UK

**Keywords:** *Marchantia polymorpha*, Enhancer Trap, Gemma, Rhizoid, Oil Cell, Margin Tissue, Meristem, Auxin, Sporeling

## Abstract

A combination of streamlined genetics, experimental tractability and relative morphological simplicity compared to vascular plants makes the liverwort *Marchantia polymorpha* an ideal model system for studying many aspects of plant biology. Here we describe a transformation vector combining a constitutive fluorescent membrane marker with a nuclear marker that is regulated by nearby enhancer elements, and use this to produce a library of enhancer trap lines for Marchantia. Screening gemmae from these lines allowed for identification and characterization of novel marker lines, including markers for rhizoids and oil cells. The library allowed the identification of a margin tissue running around the thallus edge, highlighted during thallus development. Expression of this marker is correlated with auxin levels. We generated multiple markers for the meristematic apical notch region, which have different spatial expression patterns, reappear at different times during meristem regeneration following apical notch excision and have varying responses to auxin supplementation or inhibition. This reveals that there are proximodistal substructures within the apical notch that could not be observed otherwise. We employed our markers to study Marchantia sporeling development, observing meristem emergence as defining the protonema-to-prothallus stage transition, and subsequent production of margin tissue during the prothallus stage. Exogenous auxin treatment stalls meristem emergence at the protonema stage but does not inhibit cell division, resulting in callus-like sporelings with many rhizoids, whereas pharmacologically inhibiting auxin synthesis and transport does not prevent meristem emergence. This enhancer trap system presents a useful resource for the community and will contribute to future Marchantia research.

**Significance Statement:** The liverwort *Marchantia polymorpha* is an emerging model species with huge potential for synthetic biology. We generated an enhancer trap library as a resource for Marchantia research and characterized marker lines for several important cell types (rhizoids, thallus margin, apical notch and oil cells). These marker lines can be used to track cellular dynamics in growing plants, exemplified by studying auxin responses and sporeling growth, revealing new insights into thallus development and cell fate specification.

## Introduction

The liverwort *Marchantia polymorpha* has emerged as an increasingly important model system with which to study plant evolution, development and molecular biology (Shimamura, 2016; Bowman *et al*., 2017). There are established protocols for genetic manipulation of Marchantia sporelings by Agrobacterium, which can yield thousands of transformants per transformation experiment (Ishizaki *et al*., 2016; Sugano *et al*., 2014; Sauret-Güeto *et al*., 2020). The haploid gametophyte is the dominant phase of the life cycle, which makes phenotypic characterization straightforward as recessive alleles are not a limitation. Liverworts occupy an important phylogenetic position within the land plant phylogeny, forming the bryophyte clade together with mosses and hornworts (Su *et al*., 2021; Harris *et al*., 2020; Puttick *et al*., 2018). Sequencing the Marchantia genome revealed that it is highly streamlined, with relatively low levels of genetic complexity and no evidence for whole genome duplication events, unlike mosses (Bowman *et al*., 2017; Rensing *et al*., 2007; Rensing *et al*., 2020). Liverworts have a degree of morphological complexity, with a variety of tissue and cell types (Shimamura, 2016).

The plants themselves have fast growth on defined agar media and controlled growth conditions, and can be propagated asexually by excising fragments of the thallus. Furthermore, Marchantia thalli produce clonal propagules (gemmae) which arise from a single progenitor cell within a gemma cup, meaning that clonal lines can be easily established after only a few weeks. Gemmae have a stereotypical pattern of growth and germinate almost immediately after removal from a gemma cup. Growth proceeds in an open fashion and is directly accessible for observation by live microscopy. The gemma itself is a roughly ovoid flattened sheet of cells, with a stalk scar at the former point of attachment to the gemma cup, and two apical notches approximately perpendicular to this on either side of the gemma. In the centre of the gemma there are numerous rhizoid precursors; large cells that are polygonal in shape (Shimamura, 2016). Running around the periphery of the gemma, and dotted through the central region of the thallus, are oil cells. Oil cells contain membranous organelles called oil bodies, which produce secondary metabolites that provide anti-herbivore defence (Kanazawa *et al*., 2020). The edge of the gemma is only one cell layer thick, however the gemma is thicker further towards the centre where parenchyma tissue is enclosed between two layers of epidermal tissue.

The initial phase of gemma germination sees the rapid emergence of rhizoids on both the dorsal and ventral side of the gemma. Initially the gemma has no dorsoventral asymmetry, with this axis being established after 48 hours due to gravitation cues (Bowman *et al*., 2016; Miller and Voth, 1962). After this, the ventral side features rhizoids (pegged and smooth), and mucilage papillae (Shimamura, 2016). Cell division contributes to rapid gemma growth, and occurs at the meristematic areas at the apical notches (Taren, 1958). As cells are displaced outwards from the notch region they cease division and begin expanding, eventually reaching a mature state. After 3-5 days a split appears in the thallus around the notch, forming dorsoventral ‘flaps’ of thallus. After this split, air chambers and pores begin to appear (Apostolakos and Galatis, 1985). These repeating structures are formed by stereotypical patterns of divisions, which create an open space lined by photosynthetic filaments. The chamber is connected to the external environment by an air pore bounded by elongated, curved pore cells. Oil cells are found at various locations between the air pores. After approximately 7 days, bifurcation of the apical notches produces the branched thallus morphology (Solly *et al*., 2017), and after 2-3 weeks gemma cups appear on the thallus midrib (Kato, Yasui, *et al*., 2020).

The combination of streamlined genetics, clonal growth, experimental tractability and moderate morphological complexity makes the Marchantia gemma an ideal system with which to study plant development. Gemmae have already been employed to examine processes as diverse as meristem regulation (Hirakawa *et al*., 2019; Hirakawa *et al*., 2020), auxin signalling (Eklund *et al*., 2015; Flores-Sandoval *et al*., 2015; Suzuki *et al*., 2021), ABA action (Eklund *et al*., 2018), rhizoid growth (Thamm *et al*., 2020), secondary metabolism (Takizawa *et al*., 2021), organelle dynamics (Kimura and Kodama, 2016), morphogenesis (Furuya *et al*., 2018) and light signalling (Inoue *et al*., 2019). In addition to this, Marchantia gemmae have high capacity for regeneration, providing a further method for creation and propagation of transgenic lines (Tsuboyama-Tanaka *et al*., 2015). Marchantia gemmae are a promising chassis for synthetic biology, with the aim of using it to design methods for engineering plant growth and synthesis of useful molecules. Important advances have been made in this field by the development of methods for chloroplast transformation (Frangedakis *et al*., 2021), modular modification of the Marchantia auxin signalling machinery (Kato, Mutte, *et al*., 2020), and the regulation of oil body formation (Kanazawa *et al*., 2020). A better understanding of the processes and patterns underlying gemma development would greatly boost this research. New cell and tissue specific reporters are required to highlight dynamic changes in gene expression, cell identity and specification of tissues. Marker systems developed for gemmae can also be employed for other aspects of the Marchantia life cycle, such as sexual structures, the diploid sporophyte or spore germination (Shimamura, 2016).

One powerful method for generating markers is to use an enhancer trap scheme. This involves designing a construct that fuses a reporter gene (e.g. GUS, *lacZ*, *GFP* etc.) to a minimal promoter sequence. The minimal promoter sequence typically comprises a TATA box and transcription start site, which is insufficient to drive the reporter gene expression by itself. The construct is then transformed into the study organism (e.g. by Agrobacterium mediated T-DNA insertion) and becomes incorporated at a random location within the genome. If the site of insertion is close enough to a regulatory sequence (an enhancer or promoter proximal element (Blackwood and Kadonaga, 1998)) then this drives expression of the reporter gene. The phenotype can then be examined, to see when, where and under what physiological conditions the signal appears. This basic enhancer trapping technique has been further elaborated and improved, for example by incorporating the GAL4-UAS transcriptional activator system (Haseloff, 1999; Engineer *et al*., 2005; Brand and Perrimon, 1993).

Enhancer trap systems were first developed in Drosophila (O’Kane and Gehring, 1987) and have since been used in a range of animal systems, such as mice (Shima *et al*., 2016) and zebrafish (Trinh and Fraser, 2013). In plants, however, enhancer trap approaches are especially powerful to track cell fates, given the flexibility of plant cellular identity and importance of positional cues over lineage effects (Van den Berg *et al*., 1995; Scheres, 2001; Yu *et al*., 2017). Enhancer trap libraries have been produced for several plant systems, most notably Arabidopsis (Haseloff, 1999; Radoeva *et al*., 2016; Laplaze *et al*., 2005; Campisi *et al*., 1999; Sundaresan *et al*., 1995; Klimyuk *et al*., 1995), but also rice, lotus, tomato and moss (Buzas *et al*., 2005; Pérez-Martín *et al*., 2017; Wu *et al*., 2003; Hiwatashi *et al*., 2001; Johnson *et al*., 2005). In Arabidopsis, enhancer trap methods have been particularly useful for marking cell and tissue types in systems with tightly defined growth and developmental dynamics, such as in roots (Zhang *et al*., 2019) or stomatal guard cells (Gardner *et al*., 2009) or for examining gene expression under specific physiological conditions (Baxter-Burrell, 2003). Enhancer trap systems have also yielded new mutant phenotypes and allowed the identification of novel gene functions, for example in tomato fruit development (Pérez-Martín *et al*., 2017). Enhancer trap lines can mark phenotypes that could not otherwise be easily accessible through conventional, loss-of-function genetic screens, for example, due to lethal effects of disrupting gene activity, or cryptic mutant phenotypes (Rojas-Pierce and Springer, 2003). In such situations, locating the site of insertion within the genome can identify new gene functions and important novel promoter elements, that can then be used in further research (Radoeva *et al*., 2016; Román *et al*., 2020).

Here we further extend the use of enhancer trapping to a new plant group, the liverworts, by describing the production of an enhancer trap library in Marchantiaand using it to investigate early gemma growth. Our enhancer trap construct includes a constitutive membrane marker to outline individual cells, allowing for easier identification of cell types, recognition of individual marked cells, and from this tracking of cell divisions and cell lineages. The fast growth, open development and ease of imaging fluorescent protein marker signals *in vivo* was ideally suited to the enhancer trap approach. The work produced new markers for both easily recognizable cell types (rhizoids, oil cells) and poorly understood cellular processes (specification of the meristem/notch region during gemma development, sporeling growth and thallus regeneration). It also allowed identification and characterization of a distinct tissue type, consisting of cells found running around the edge of the gemma. These markers were useful to study other developmental processes, such as spore germination and growth responses to auxin treatment. The establishment of this library in Marchantia provides new resources for future research.

## Methods

### Vector Construction and Transformation

The enhancer trap construct was built according to the Loop assembly protocol (Pollak *et al*., 2019; Sauret-Güeto *et al*., 2020) from L0 parts, L1 and L2 plasmids in the OpenPlant Loop assembly toolkit (Sauret-Güeto *et al*., 2020). The annotated plasmid sequences of L2_GAL-UAS_mV-N7-eG-Lt-CSA (L2_238-CsA, p5-35Sx2:HygR p5-UAS:mVenus-N7 p5UBE2:eG-Lt del35:GAL4-VP16) (=ET238) and L2_GAL-UAS_mV-N7_mS-Lt-CSA (L2_239-CsA, p5-35Sx2:HygR p5-UAS:mVenus-N7 p5UBE2:mS-Lt del35:GAL4-VP16) (=ET239) are given in Appendix S1. L2_GAL-UAS_mV-N7_eG-Lt-CSA (L2_238-CsA, p5-35Sx2:HygR p5-UAS:mVenus-N7 p5UBE2:eG-Lt del35:GAL4-VP16) was done by combining pCsA, L1_HygR-Ck1, L1_UBE2:eG-Lt-Ck3 from the OpenPlant toolkit and new L1 plasmids L1_UAS:mV-N7-Ck2 and L1_del35:GAL4-VP16_rev-CK4. L1_UAS:mV-N7-Ck2 (p5-UAS mVenus-N7) was done with OpenPlant toolkit parts: plasmid pCK2, CD12_mVenus, CTAG_linker-N7 and 3TERM_Nos-35S and new synthesised L0 part PROM5_UASGAL4_min35S. L1_GAL4_rev-CK4 (p5-del35:GAL4-VP16_rev-CK4) was done with new synthesised L0 parts 3TERM_Nos_rev and PROM5CDS_del35S:GAL4-VP16_rev and plasmid pCK4 from OpenPlant toolkit. L2_GAL-UAS_mV-N7_mS-Lt-CSA (L2_239-CsA, p5-35Sx2:HygR p5-UAS:mVenus-N7 p5UBE2:mS-Lt del35:GAL4-VP16) was done by combining pCsA, L1_HygR-Ck1, L1_UAS:mV-N7-Ck2, L1_UBE2:mS-Lt-Ck3 and L1_del35:GAL4-VP16_rev-CK4. The minimal TATA box promoter, GAL4-VP16 and mVenus-N7 sequences were positioned immediately adjacent to the right border, which is first integrated into the plant genome, and provides the best opportunity for precise integration of critical elements of the enhancer trap element.

MpPIN and MpYUC2 marker lines were also constructed by Loop assembly to produce L2_268 35SBP:HygR2, p5-PIN1:GAL4-VP16, p5-UAS-GAL4:mVen-N7, o941:eGFP-Lti6b and L2_295 35S:HygR, p5-ERF20:mVen-N7, p5-YUC2(5.5kb):mTurq-N7, p5-o941:mScar-Lti6b. Full annotated plasmid sequences are provided in Appendix S1. The 5’ promoter sequence for MpPIN1 started at TGAATGCGGTCGAGGACGGG (2.977kb upstream of the start codon). The 5’ promoter sequence for MpYUC2 started at GTGGTTCCGCTCTCTCGTCG (4.97kb upstream of the start codon).

The plasmid vectors were transformed into Marchantia sporelings (Cam accession) by Agrobacterium-mediated transformation according to the method described in Ishizaki *et al*., 2016; Sauret-Güeto *et al*., 2020. Transformed sporelings were grown for 2 weeks at 21°C under continuous light (intensity= 150 μmol/m^2^/s) on 1.2% w/v agar (Melford capsules A20021) plates of Gamborg’s B5 media with vitamins (Duchefa Biochemie G0210) prepared at half the manufacturer’s recommended concentration and adjusted to pH 5.8. The media was supplemented with 100µg/ml cefotaxime (BIC0111; Apollo Scientific, Bredbury, UK) to cure bacteria growth, and 20µg/ml hygromycin (10687010; Invitrogen) for plant selection. This media and antibiotic concentrations were used for all further growth experiments unless stated otherwise.

### Enhancer Trap Screening, Culturing and Characterization

A limited preliminary transformation and screening experiment was carried out to test the enhancer trap vector constructs, before a full round of transformation and screening was performed. In the full screen, 101 transformants from the plasmid ET238 transformation were selected and isolated for further characterisation. 353 transformants from the plasmid ET239 transformation were selected at random and transferred to fresh plates for further characterisation. In addition, the remaining transformants were screened for mVenus fluorescence using a Leica M205 FA stereomicroscope. Fluorescing sporelings were identified, isolated and removed for further growth on fresh plates. Each sporeling was named as ET238-PX or ET239-PX, where PX denotes the individual plant line, numbered in order of isolation.

T0 plants were monitored for the presence and distribution of fluorescent signal as the plants grew. Gemmae were taken only from gemma cups located in regions of the thallus with signal, since the clonal origin of the gemma (Kato, Yasui, *et al*., 2020) avoids problems arising due to chimeric plants. 3-6 gemmae from each transformant were removed from the gemma cups and grown on ½ Gamborg’s media agar plates in the presence of hygromycin and cefotaxime. Gemmae were examined from one day after removal from the gemma cup (i.e., 1 day post-germination, 1dpg) until 7dpg using a Leica M205 FA stereomicroscope equipped with filters Plant RFP (excitation filter ET560/40 nm, emission filter ET594/10 nm), ET GFP (ET470/40 nm, ET525/50 nm), ET YFP (ET500/20 nm, ET535/30 nm), ET Chlorophyll LP (ET480/40 nm, ET610 nm LP), and ET GFP LP500 (ET470/40 nm, ET500 nm LP), used respectively for observing mScarlet-I, eGFP, mVenus, chlorophyll, and eGFP together with chlorophyll autofluorescence (Sauret-Güeto *et al*., 2020). Patterns of fluorescence (nuclear mVenus and membrane eGFP/mScarlet) were noted. Fluorescence was categorized into broad patterns according to the tissue or cell type involved. If no fluorescence was observed on any gemma by 7dpg the line was discarded. G1 lines exhibiting gemma fluorescence were maintained on selective media by subculturing or by propagation of gemmae and grown in a Panasonic RLR-352 PE Climate Chamber at 16°C under continuous light at intensity below 100 μmol/m^2^/s. Lines were preserved as gemmae at 4°C on agar (Sauret-Güeto *et al*., 2020) and preserved in long-term frozen storage at -80°C using the CRUNC method (Takahashi and Kodama, 2020).

Plant lines with strong and consistent patterns of fluorescence that marked specific regions, tissues or cells were selected for further characterization. These lines were maintained in Aralab Fitoclima 600 growth cabinets at 21°C under constant light for faster growth. Gemmae were grown on 50mm agar plates containing ½ Gamborg’s media with hygromycin and cefotaxime for imaging using a Leica SP8 confocal microscope. Images were taken daily between 0dpg and 4dpg. 4-5 gemmae were imaged repeatedly each day with a HC PL APO 20x/0.75 CS2 air objective; three other gemma were mounted on slides for imaging using a HC PL APO 40x/1.30 CS2 oil objective. Excitation and collection settings for each fluorophore are given in Table S1. Images were taken in photon counting mode with sequential scanning. Time gating was active to supress autofluorescence. Maximum-intensity projections of the images were obtained from z-stack series, capturing z-slices at intervals between 1μm-12.5μm. Confocal microscope images are shown with mVenus channel in yellow, mScarlet channel in red, chlorophyll autofluorescence in grey, unless otherwise noted. Stereomicroscope images were taken with a Leica M205 FA stereomicroscope equipped with a Leica DFC465 FX camera using the Plant RFP filter (excitation filter ET560/40nm, emission filter ET594/10 nm) for the mScarlet channel, ET YFP filter (excitation filter ET500/20nm, emission filter ET535/30nm) for the mVenus channel and ET Chlorophyll LP filter (excitation filter ET480/40nm, emission filter ET610nm LP) for the chlorophyll autofluorescence channel. Images were taken with the 2x/0.10 objective at variable zoom according to the size of the gemma being imaged. Exposure time for the chlorophyll channel was 5ms, 9 gain; for the mVenus channel 3000ms, 1.7gain; and for the mScarlet channel 100ms, 9 gain. Gamma was set at 0.9. Z-stack series were taken at intervals between 184µm and 1.18mm (for control plants) according to the size of the gemma being imaged. Extended depth of field images were created from Z-stacks with the Leica LasX software package, using the mVenus channel as reference. All images were processed using the Leica LasX and Fiji/ImageJ software packages. Figures were created using the Stitching (Preibisch *et al*., 2009) and QuickFigures (Mazo, 2021) plugins. Gemma images are presented with the stalk scar orientated towards the bottom and apical notches approximately perpendicular at each side, unless otherwise noted. Excised gemma fragment and sporeling images are presented at various orientations, with apical notches marked where relevant.

### Spore Production

Six lines were selected for spore production by crossing with wild-type Cam-1 (male) or Cam-2 (female) plants. Plants were grown in Sac O2 Microboxes for crossing and spore production, as described in Sauret-Güeto *et al*., 2020. Spores were stored at -80°C and sterilized before planting using the method described in (Sauret-Güeto *et al*., 2020). Time course microscopy experiments were performed using spores germinated and grown on 50mm agar plates containing ½ Gamborg’s media with hygromycin and cefotaxime in the same growth cabinets and conditions used for gemmae growth. Spores and sporelings were imaged under the same conditions as for gemmae.

### Identification of the Genomic Location of Insert Site

gDNA was extracted from wild-type crossed sporelings grown under hygromycin selection. Sporelings were harvested after 4.5 weeks growth and the tissues preserved at - 80°C. gDNA extraction was performed using a modified CTAB-based protocol (Frangedakis, 2019). Insert sites were identified using a thermal asymmetric interlaced PCR based method (TAIL-PCR) (Liu *et al*., 1995) based on Grotewold, 2003. Primer sequences, PCR mixes and PCR protocols are given in Methods S1. PCR products were run on an agarose electrophoresis gel and the strongest bands excised for purification using a QIAquick Gel Extraction Kit (Qiagen). The purified PCR products were sequenced directly using Sanger sequencing by Genewiz. The nucleotide sequence generated was used to search the *Marchantia polymorpha* genome v3.1 and MpTak1v6.1 using BLAST (Altschul *et al*., 1997) on the MarpolBase (marchantia.info) and Phytozome (phytozome-next.jgi.doe.gov) databases. The insertion site locations identified were supported by the sequencing of multiple TAIL-PCR products, each from different primer combinations, that all matched to the same location in the Marchantia genome. Potential candidate genes near the insertion site were explored further using recently-released single cell RNA sequencing (scRNA-seq) data, available via wanglab.sippe.ac.cn/Marchantia-census (Wang *et al*., 2023).

### Laser Ablation

Plant tissues were cut by laser ablation using a Leica LMD6000 laser dissection microscope equipped with a solid state 355nm cutting laser. For excising apical notches, a 90µm diameter circle was drawn with its centre point at the apical notch, and all tissue within this circle was ablated. Laser settings were 60 power, 45 aperture, 25 speed. All cuts were done under the 10x objective lens. Gemmae were removed from the gemma cup (i.e. 0dpg) and planted on 50mm agar plates. Laser ablation was performed immediately, directly on the plates, and then gemma imaged using a Leica SP8 confocal microscope. Subsequent imaging was conducted daily from 0 days after cutting (DaC) until 4DaC for all lines and was continued until 5DaC in those lines where the apical notch/meristem marker had not reappeared at 4DaC. For all lines an uncut gemma, taken from the same gemma cup and grown on the same plate, was also imaged from 0dpg until 4dpg as a control. For laser ablation studies of enhancer trap line ET239-P64, gemma were cut using the same laser settings for bulk tissue cuts under the 10x objective lens, and 60 power, 35 aperture, 20 speed under the 40x objective lens for fine scale cuts or ablations of a few cells at the edge of the gemma or around the apical notch. To isolate small clusters/single cells, a trace was drawn across the gemma in pattern of radial spokes and concentric circles, and ablations done across this pattern at 60 power, 45 aperture, 30 speed under the 10x objective lens. The remaining portions of the gemma were left intact to grow and observed again at 4DaC, upon which the largest surviving pieces of tissue were destroyed by further laser ablation, in order to provide space for the single cells/clusters of cells to grow and be accessible for imaging. This process was repeated where necessary during the time course until the single cells/clusters of cells of interest had fully regenerated into thalli.

### Auxin Manipulation

For growing gemmae with elevated levels of auxin, 50mm agar plates containing ½ Gamborg’s media with hygromycin and cefotaxime were prepared with addition of 1µM, 3µM or 5µM 1-naphthaleneacetic acid (NAA) (Sigma) dissolved in 1M NaOH and then diluted in water. Auxin synthesis inhibitor treatments were done by growing gemmae on plates made with 10µM 5-(4-chlorophenyl)-4H-1,2,4-triazole-3-thiol (yucasin) (ChemCruz, Santa Cruz Biotechnology) or 100µM L-kynurenine (Sigma) dissolved in DMSO (Tsugafune *et al*., 2017). Auxin transporter inhibitor treatments were done by growing gemmae on plates made with 100µM 2,3,5-triiodobenzoic acid (TIBA) (Aldrich) dissolved in water. Media plates were stored at 4°C wrapped in foil to protect from light and were used within 6 weeks of preparation. Auxin treatment experiments were all conducted with plants grown in the same growth cabinets as the control plants, with 4-7 gemmae grown under each treatment.

## Results and Discussion

### Generating and Screening the Marchantia Enhancer Trap Library

Sporelings were transformed with either the ET238 or ET239 enhancer trap vector, following an established protocol (Sauret-Güeto *et al*., 2020) and then selected on media containing hygromycin. Sporelings were picked out at random and these T0 plants grown until gemma cups formed. Gemmae were isolated, germinated and the G0 plants monitored for nuclear mVenus signal appearing in the first 7 days.

The full screen involved transformation of sporelings with either the ET238 or ET239 vector, as both constructs showed equally promising results in pilot experiments. Having alternative constructs will allow for flexibility in reducing potential fluorescence emission spectra cross-talk in downstream imaging experiments or for crossing enhancer trap lines with lines containing other fluorescent markers. Equal volumes (150µl per transformation) of starting spore suspension taken from the same pooled sterilized sporangia extracts were used for each construct for the full screen. After 10 days, transformed sporelings were selected at random, picked out, and two days later were observed by stereomicroscope to identify sporelings with observable fluorescent reporter signals (mVenus or eGFP/mScarlet as appropriate). These T0 plants were isolated for further analysis. In total 101 ET238-construct sporelings and 353 ET239-construct sporelings were taken to this stage. Isolated sporelings were then monitored for fluorescent reporter signals; plants with fluorescent signal were retained and the general distribution of signal noted (e.g. widespread, partial/chimeric, etc.); those plants with no observable signal were disposed of. It should be noted that plants without signal may not simply be due to transformation failures or silencing; it is possible that fluorescent signal would be observed under different growth conditions or at different developmental stages than T0 sporelings of this age. After this stage 49 ET238 plants and 190 ET239 plants remained.

Approximately 2-3 weeks after isolation gemma cups appeared on these T0 plants. Where there were indications of chimeric transformants in the T0 generation, gemmae (G0) were only taken for screening from gemma cups that developed within parts of the plant displaying consistent signal. G0 gemmae were screened across seven days. Only lines with nuclear mVenus reporter signal observed within this timeframe were retained after this screening phase: 32 ET238 lines and 125 ET239 lines. These lines were maintained in culture with some lines subsequently disposed of due to problems with fungal contamination or loss of expression due to gene silencing. In total, 26 ET238 lines were cultured with 25 cryopreserved, and 122 ET239 lines were cultured with 121 cryopreserved. The statistics for each screening phase are given in Table 1. The percentages of selected plant lines that display reporter signal are broadly in line with enhancer trap projects from other plant species, such as Arabidopsis (48%), rice (26-59%), tomato (32-43%) and moss (30%) (Sundaresan *et al*., 1995; Pérez-Martín *et al*., 2017; Hiwatashi *et al*., 2001; Wu *et al*., 2003).

**Table 1.** Statistics on the number of lines at each screening step. Numbers are expressed as a percentage of sporelings isolated or isolated sporelings with signal (number in brackets). All observations were made by fluorescence stereomicroscope and “mVenus signal” refers to the presence of consistent nuclear reporter expression that was observable in multiple gemmae replicates.

| Screening Stage | ET238 | % ET238 | ET239 | % ET239 |
| --- | --- | --- | --- | --- |
| Sporelings isolated | 101 | 100% | 353 | 100% |
| Isolated sporelings with mVenus signal | 49 | 49%<br>(100%) | 190 | 54%<br>(100%) |
| G0 gemma with mVenus signal by 7dpg | 32 | 32% (63%) | 125 | 35%<br>(66%) |
| Plants currently in culture or cryopreserved | 26 | 26%<br>(53%) | 121 | 34%<br>(64%) |

mVenus expression was observed through stereomicroscope screening of gemma one day after removal from the gemma cup (i.e., 1dpg) until seven days after germination (7dpg) in the lines that were maintained in culture (see Table 2). These signal patterns were recorded from multiple replicates from each line. Signal types were categorized according to mVenus expression in certain structures, cell types or spatial localization, and if this association was consistent across replicates. Within these broad categories of expression pattern, there did exist minor variability between gemmae from the same line, or occasional degree stochastic expression in individual cells. This is possibly due to leaky expression, cryptic variation in growth conditions, the typical morphological variability in Marchantia and the general complexity of enhancer activity across different cell and tissue types in multicellular organisms (Jenett *et al*., 2012). The signal patterns listed in Table 2 were subject to detection biases in favour of more obvious localization patterns (e.g. in the notch region), signal appearing in more easily recognizable cell types (e.g. oil cells, rhizoids), and in more prevalent or prominent morphological features (e.g. air chambers, distribution across the gemma centre or margins).

**Table 2.** Numbers of lines with expression pattern in selected morphological features. Expression pattern means the areas or cell types where nuclear mVenus signal was observed by stereomicroscope in the first week after germination. The listed patterns had strong expression, were exclusive to or clearly concentrated in the indicated feature, were consistent across multiple replicate gemmae and persisted through the entire observation period. Other lines are present in the enhancer trap library that mark other morphological features; or have expression patterns in these morphological features but with weaker expression or non-exclusive/concentrated distribution.

| Expression pattern in 0dpg-7dpg gemma | Number of lines |
| --- | --- |
| Meristem/apical notch | 30 |
| Oil cells | 5 |
| Rhizoid-specific | 13 |
| Ventral surface but not rhizoids | 1 |
| Air chambers and air pores | 38 |
| Thallus periphery | 2 |
| Central zone | 3 |
| Absent from central zone | 3 |
| Subepidermal tissues | 2 |
| Bulge behind apical notch | 2 |
| Widespread/ubiquitous expression | 11 |

Notwithstanding this, we selected some examples to exemplify the utility of the enhancer trap approach in Marchantia. These lines had clear, reliable signals that appeared consistently and closely associated with certain cell types or thallus regions, with small amounts of variation between individual gemma. These lines were characterized for growing gemmae from 0dpg to 3dpg and then used for further study, such as molecular characterization or response to phytohormone treatments.

### Rhizoids

Numerous enhancer trap lines showed nuclear mVenus signal in the dorsal rhizoids and/or rhizoid precursor cells. These cell types are easily recognizable, are present early in gemma development and contain large, prominent nuclei, making fluorescence relatively easy to observe. Table 3 lists the various types of rhizoid signals that were observed. Seven enhancer trap marker lines were specific to rhizoid-type cells present in all rhizoids (dorsal and ventral), while a further six lines had signal specific to rhizoid-type cells and the ventral surface of older gemma. It is noteworthy that no lines were found with dorsal rhizoid-specific expression, supporting the notion that the dorsal rhizoids of early germinating gemma are not a unique cell type and that the gemma initially lacks fully differentiated dorso-ventral asymmetry (Bowman *et al*., 2016; Miller and Voth, 1962). Early rhizoid signals were often seen in addition to nuclear signals being observed in other cells in the thallus, but this did not persist in the older thalli where only ventral rhizoids were growing, and mVenus marker expression was often irregularly distributed in some but not all rhizoids in the early gemma.

**Table 3.**
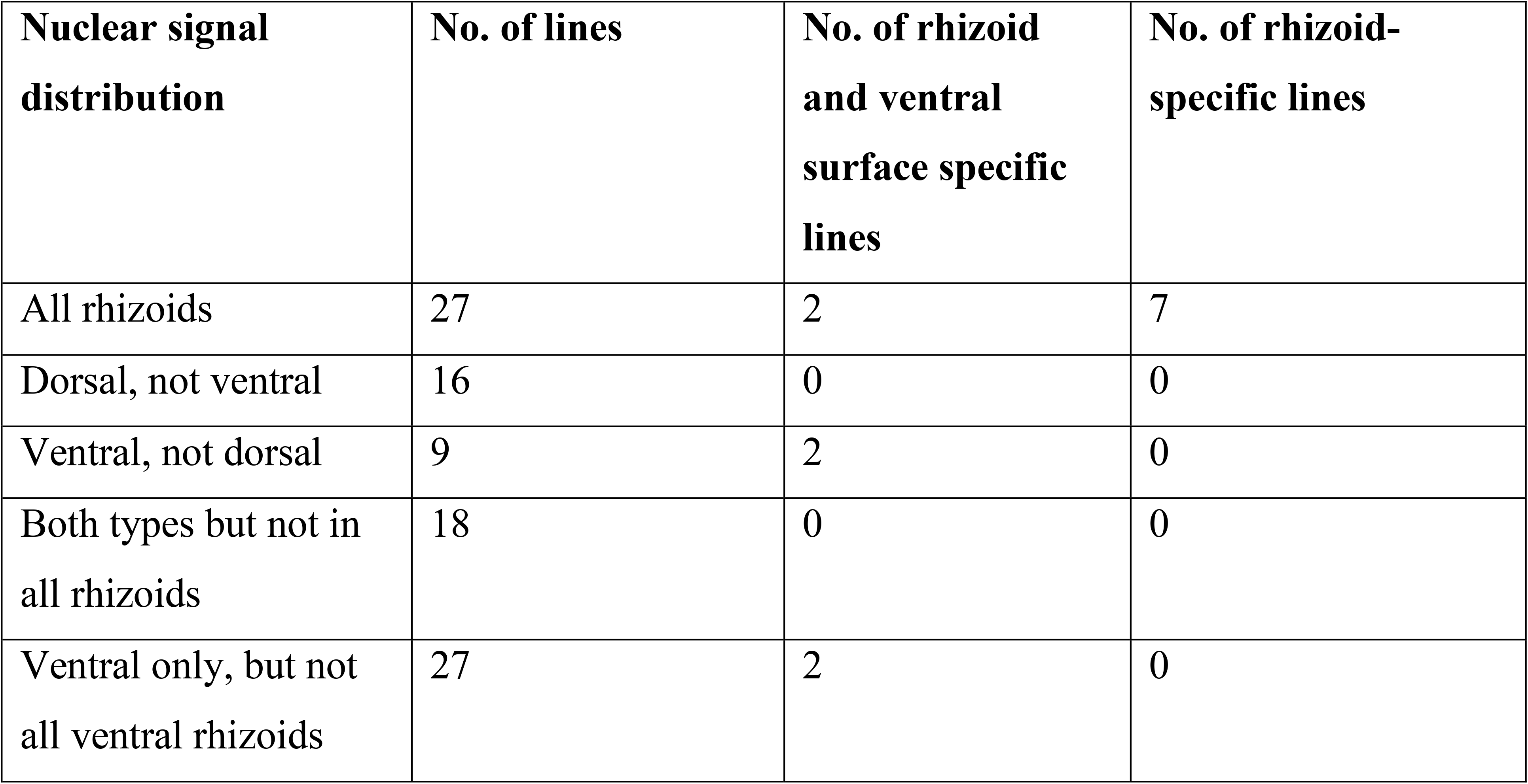
Numbers of lines with various rhizoid-related reporter signal. Marker expression was observed by stereomicroscope in gemmae up to 7dpg, lines were included if rhizoid signal was observed in some but not all of this period. For the 13 lines with rhizoid specific or rhizoid and ventral surface specific expression, fluorescent patterns were observed across multiple replicates and expression was persistent through the observation period. The ventral rhizoid classification includes both smooth and pegged rhizoid types.

Line ET239-P177 was selected as a representative rhizoid-specific marker line. It marked all rhizoid types: dorsal and ventral, pegged and smooth, as well as rhizoid precursor cells in the 0dpg gemmae, but not non-rhizoid cells on the ventral surface of older gemmae. Confocal scanning microscope images of the marked rhizoids in this line are shown in Figure 1 (a)-(d). The insertion site in line ET239-P177 was identified as being at an intergenic site on chromosome 3 (18170559 in the MpTakv6.1 genome annotation). A putative candidate gene near this site is Mp3g17930 (Mapoly0039s0003) (Figure 1(e)), annotated in the *Marchantia polymorpha* v6.1 genome as being the Marchantia homolog of *Root Hairless-like* 6. This is a transcription factor previously identified as being crucial for rhizoid development in liverworts and root development in vascular plants (Proust *et al*., 2016). This would tally with the observations of rhizoid marker signal in line ET239-P177, notwithstanding the uncertainties related to identifying the gene whose enhancer element has been trapped. Mp3g17930 does not appear in any of the identified scRNA-seq data, but its UMAP plot does show some expression within the rhizoid cluster (Wang *et al*., 2023). ET239-P177 therefore represents a line from this enhancer trap library marking a known cell type with obvious morphology, and likely linked to a gene previously known to be related to the development of that cell type.

**Figure 1.**
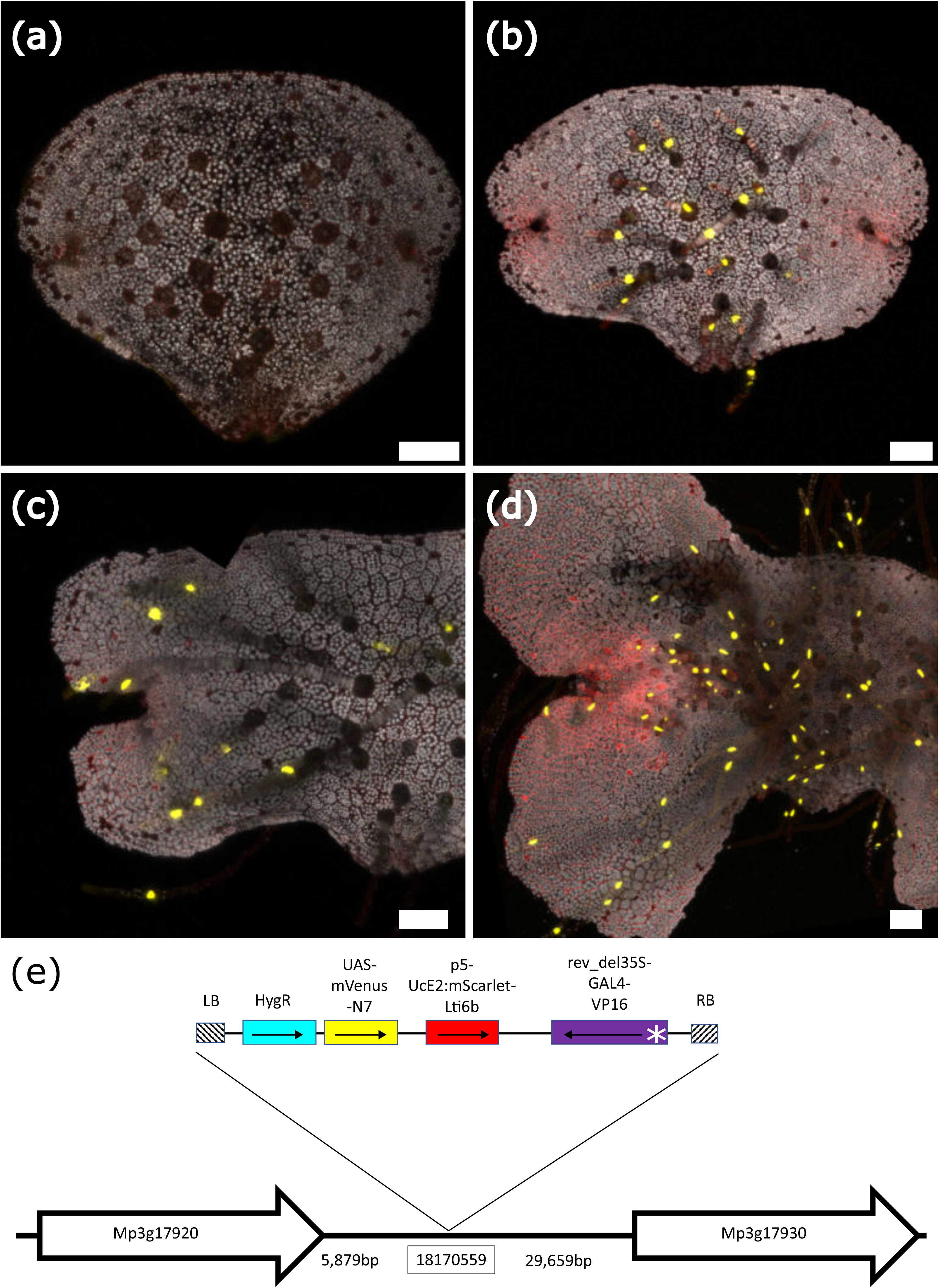
Rhizoid marker line ET239-P177. The same gemma imaged on 0dpg (a), 2dpg (b) and 3dpg (c), showing localization of the mVenus marker signal to the rhizoid cells. (d) shows the ventral side of a 6dpg gemma, demonstrating that the marker signal is unique to rhizoids and is present in all rhizoid types. Scale bars= 100µm. (e) schematic illustrating the genomic location of the insert site on chromosome 3. A possible candidate gene nearby is Mp3g17930 (Mapoly0039s0003), the Marchantia homolog of the root-associated transcription factor RSL6. Neighbouring genes are indicated as white arrows, starting at the gene’s start codon and pointing towards the gene’s stop codon. Schematics are not drawn to scale and distances are indicated in base pairs. The T-DNA map shows the relative positions of the minimal promoter (white asterisk), GAL4 enhancer (purple), mScarlet membrane marker (red), UAS-mVenus nuclear marker (yellow) and hygromycin resistance gene (cyan), as well as the left and right border sequences (hatched). Black arrows point from the gene’s start codon towards the stop codon.

### Oil Cells

Oil cells are a cell type containing oil bodies (a membrane-rich organelle containing various secondary metabolites including terpenoids and (bis-)bibenzyl compounds) found throughout the Marchantia thallus (Kanazawa *et al*., 2020). Identification of oil cells depends on recognizing these oil bodies (clearly distinguishable with a membrane marker, Figure 2(a)) and on other features such as having fewer and smaller chloroplasts. In the 0dpg gemma, oil cells are found at regular intervals around the circumference of the gemma, set inwards from the gemma edge, and are also dispersed throughout the gemma body. As the plant grows, oil cells continue to be sited close to the edge of the thallus but are also dispersed throughout the thallus, including between air chambers. In this case, identification of oil cells becomes more complicated due to their irregular distribution.

**Figure 2.**
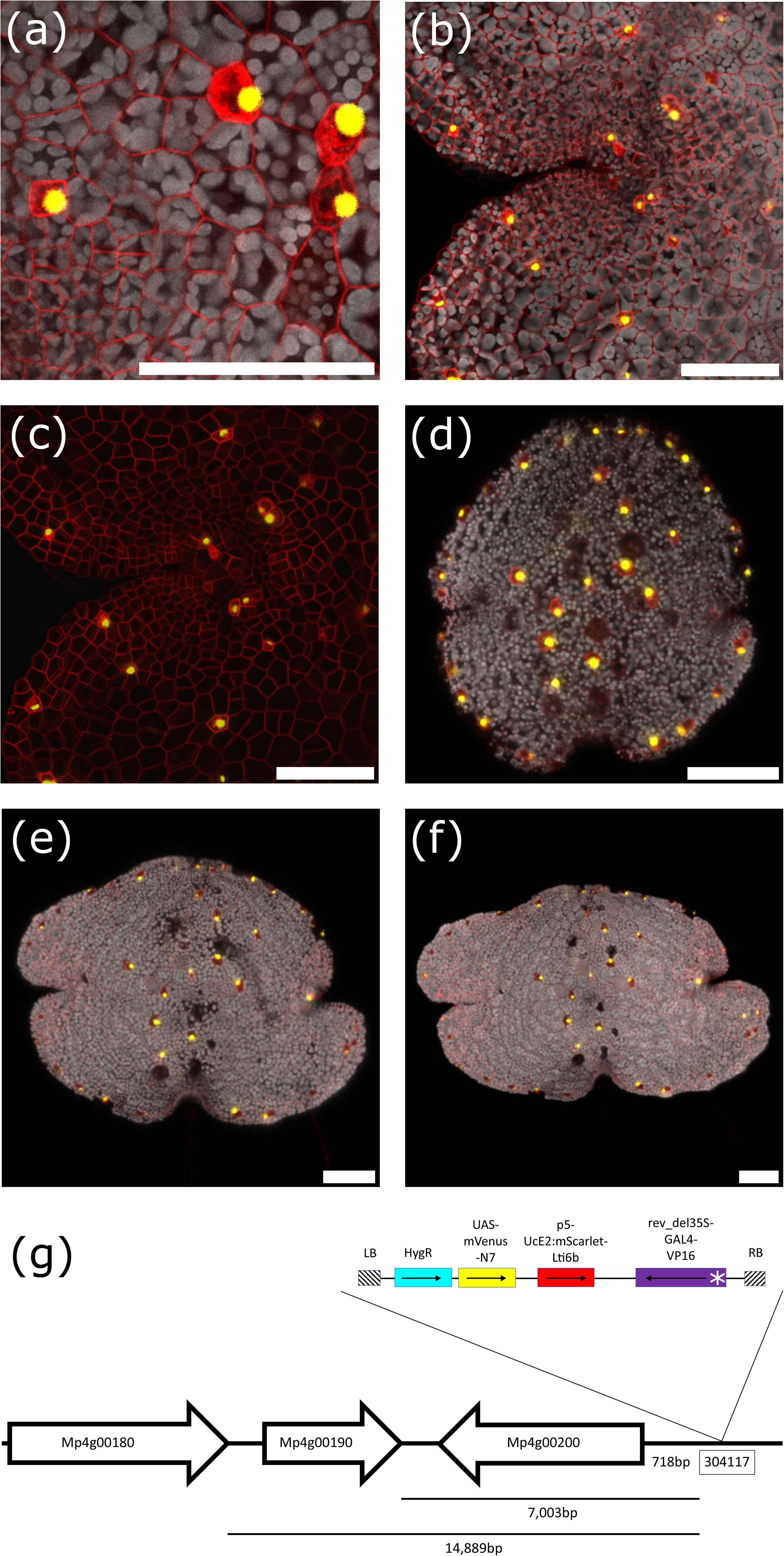
Oil cell marker line ET239-P54. (a) The mScarlet membrane marker highlights the dense membranes of the oil bodies. (b) and (c) show a close up of a portion of a 3dpg gemma with (b) and without chlorophyll channel (c), demonstrating that the marker signal in ET239-P54 is unique to the oil cells. The same gemma imaged at 0dpg (d), 2dpg (e) and 3dpg (f), showing localization of the mVenus marker signal to the oil cells. Scale bars = 100µm. (g) illustrates the genomic location of the insert site on chromosome 4, near possible candidate WRKY family transcription factor genes Mp4g00200 (Mapoly0162s0001) and Mp4g00180 (Mapoly0162s0003). Neighbouring genes are indicated as white arrows, starting at the gene’s start codon and pointing towards the gene’s stop codon. Schematics are not drawn to scale and distances are indicated in base pairs. The T-DNA map shows the relative positions of the minimal promoter (white asterisk), GAL4 enhancer (purple), mScarlet membrane marker (red), UAS-mVenus nuclear marker (yellow) and hygromycin resistance gene (cyan), as well as the left and right border sequences (hatched). Black arrows point from the gene’s start codon towards the stop codon.

Five marker lines were identified with signals specific to oil cells. One line, ET239-P54, was chosen as a representative oil cell line due to its specificity and strong, reliable signal (Figure 2(b)-(f)). The formation of new oil cells during thallus growth was also observable around the apical notch region, indicating the zone in which oil cell fate specification occurs (Kanazawa *et al*., 2020) and where the spatial dynamics controlling the distribution of oil cells happens. The insertion site in line ET239-P54 was identified as being on chromosome 4, at site 304117 in the MpTakv6.1 genome annotation. This location is close to two possible candidate genes (Mp4g00200 (Mapoly0162s0001) and Mp4g00180 (Mapoly0162s0003)) annotated in the *Marchantia polymorpha* v6.1 genome as belonging to the WRKY transcription factor family (WRKY12 and WRKY13 respectively), and a third possible candidate gene (Mp4g00190 (Mapoly0162s0002)) that encodes a small protein of unknown function and with no known homology to other plant genes (Figure 2(g)). Of these potential candidates, the WRKY12 gene Mp4g00200 is enriched within the oil cell cluster identified by scRNA-seq (Wang *et al*., 2023). WRKY transcription factors are known to be involved in abiotic stress and anti-herbivore responses (Bakshi and Oelmüller, 2014; Wani *et al*., 2021), and oil cells are known to have a role in deterring herbivores (Kanazawa *et al*., 2020). The identification and characterization of a new oil cell marker opens avenues towards producing synthetic promoters specific to these cells. This has biotechnological potential for harnessing Marchantia oil bodies for production of specific secondary metabolites such as medicinal isoprenoids (Takizawa *et al*., 2021; Suire *et al*., 2000).

### Margin Tissue

One of the most powerful aspects of an enhancer trap screen is in its ability to mark new types of cells or tissues for further study (Shima *et al*., 2016; Sundaresan *et al*., 1995). An example of this is line ET239-P64, which exhibits nuclear mVenus expression in the two rows of cells that run around the edge of the gemma, except at the stalk scar (Figure 3(a)). The signal in these marginal cells appears just beyond the apical notch, approximately 10-12 cells out from the apex cell. Here the enhancer trap line ET239-P64 allows characterization of these cells as having a specific “margin tissue” identity.

**Figure 3.**
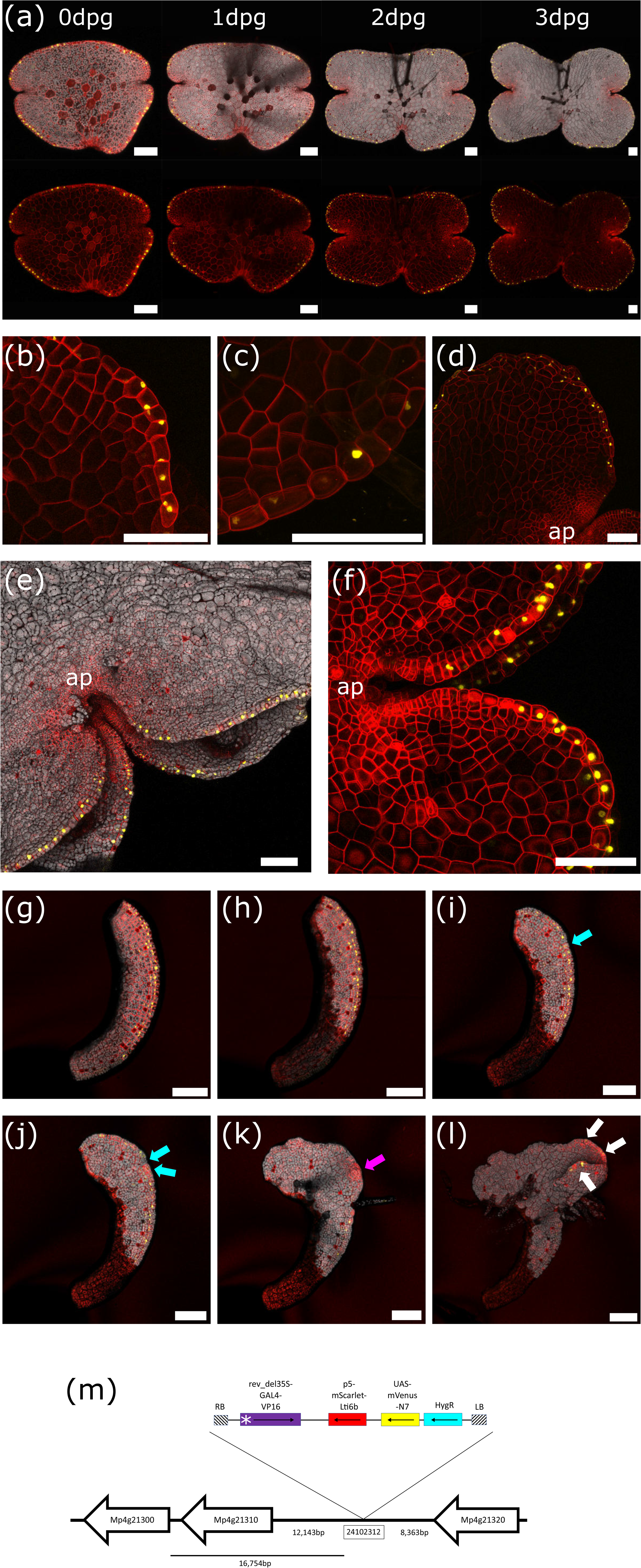
Margin tissue marker line ET239-P64. (a) The same gemma imaged at 0dpg, 1dpg, 2dpg and 3dpg, showing localization of the mVenus marker signal to the rows of cells running around the edge of the gemma, distal to the oil cells. The bottom image shows the same image with the chlorophyll channel omitted. (b) and (c) are close-up views of margin tissue cells, showing their pillow-shaped morphology. (d) Ventral view of a gemma imaged at 3dpg, indicating that the marker signal is only localized to the cells at the gemma edge, and that this is visible from both the dorsal and ventral views because this region is only one cell layer thick. “ap” indicates the location of the apical notch. (e) and (f) show the second row of marker signal that emerges as the gemma splits along its z-axis approximately 3dpg-5dpg. “ap” indicates the location of the apical notch. (g)-(l) Isolated portion of margin imaged at 0DaC (g), 1DaC (h), 2DaC (i), 3DaC (j), 5DaC (k) and 9DaC (l) showing loss of margin marker signal (cyan arrows), reprogramming into new meristem (pink arrow), before appearance of new margin tissue signal de novo (white arrows). Scale bars = 100µm. (m) is a schematic diagram illustrating the genomic location of the insert site on chromosome 4, near to a putative candidate Myb transcription factor gene Mp4g21300 (Mapoly0090s0091), annotated as Mp1R-MYB17. Neighbouring genes are indicated as white arrows, starting at the gene’s start codon and pointing towards the gene’s stop codon. Schematics are not drawn to scale and distances are indicated in base pairs. The T-DNA map shows the relative positions of the minimal promoter (white asterisk), GAL4 enhancer (purple), mScarlet membrane marker (red), UAS-mVenus nuclear marker (yellow) and hygromycin resistance gene (cyan), as well as the left and right border sequences (hatched). Black arrows point from the start codon towards the stop codon for each gene.

These cells are elongate and ‘pillow-shaped’, are found distal to a row of oil cells, and are always found as one-cell thick layers (Figure 3 (b)-(d)). This characteristic cell morphology has been previously noted as being present at key stages in early gemma development (Kato, Yasui, *et al*., 2020; Shimamura, 2016) and sporeling development (O’Hanlon, 1926). The margin tissue cells at the edge of the gemma were observed to divide both parallel and perpendicular to the gemma edge, occasionally dividing in both directions successively. While division was usually orientated across the long axis of the cell, the direction of margin tissue cell division could not be predicted from cell shape alone, and neither cell size nor morphology was an indicator of which cells would divide. No divisions occurring in the z-axis were recorded. This maintains the margin tissue as a one-cell thick layer, and as a rim of one or two cells distal to the oil cells. In common with other tissue types, cell divisions in the margins are restricted to the area surrounding the apical notch, and outside this region margin tissue cells are mitotically inactive (see Figure S1 for examples of time course imaging with margin tissue cell divisions tracked). mVenus expression is seen stronger distal from the notch, possibly indicating a gradual cell fate transition from apical notch (i.e., meristematic) cell to margin tissue cell.

At some point after the gemma is removed from the gemma cup and growth commences, a new, additional set of margin tissue forms further in, distal from the apical notch, running parallel to the notch sides (Figure 3 (e), (f); Movie S1, Movie S2). Nuclear localised mVenus expression indicates that the formation of this second set occurs by 5dpg, though in some cases it was observable as early as 1dpg. As cell division continues, and cells between the two sets of margin tissue enlarge as they move outwards from the notch region (Solly *et al*., 2017), a split forms in the thallus along the z-axis. This z-axis split marks the transition to the mature thallus form, with the appearance of air pores and air chambers in the region between the two sets of margin tissue (Apostolakos and Galatis, 1985). After this split, no new sets of margin tissues develop, and instead the existing margin tissues grow from the apical notches. This indicates that the formation of a second set of margin tissue is an early step of the developmental sequence that creates the thicker thalloid form and that the regulation of margin tissue formation is a component of the correct development of the Marchantia thallus.

Cutting experiments were conducted using a laser ablation microscope to test whether margin tissue is specified by the simple presence of a dorso-ventral boundary, or if margin tissue cell fate is determined by other cell associations during development. Various cuts were made by laser ablation to 0dpg gemmae, along the gemma edge and to the apical notch region (Figure S2(a)-(e)). After observing gemma growth at regular intervals, it was consistently observed that margin tissue mVenus signal was not simply a result of there being a dorso-ventral boundary. Only intact apical notches produced new margin tissue, as denoted by the emergence of new mVenus signal. Margin tissue signal did not appear spontaneously at the gemma edge, but instead only appeared in cells in the region immediately outside of the apical notch and was then maintained as the gemma grew larger. Crucially, if a small portion of gemma was excised from the peripheral region, margin tissue signal was only present in those cells that were at the edge of the original intact gemma (Figure 3 (g), (h)). As meristem regeneration occurs (Kubota *et al*., 2013), the margin tissue signal fades (i.e., those cells lose margin tissue specification, blue arrows, Figure 3 (i), (j)) and the cells acquire a new fate, becoming meristematic cells. This is consistent with de-differentiation of Marchantia cells during thallus regeneration (Nishihama *et al*., 2015). As the regeneration of meristematic tissues continues, a new apical notch region forms (pink arrows, Figure 3 (k)). New margin tissue is produced *de novo* from this new notch region (white arrows, Figure 3 (l)). The formation of new margin tissue occurs along the axis of the new apical notch, irrespective of other environmental cues such as gravity or light. This confirms that margin tissue is a specific tissue cell type produced from the apical notch meristem, rather than appearing purely due to external positional cues or a cell finding itself with no neighbouring cells on one side (i.e., at the edge of a gemma). This is in contrast to some cell types in the *Arabidopsis* root, where removal of one cell type causes other cells to acquire new fates by positional cues only (Van den Berg *et al*., 1995), or in the leaves of angiosperms, where the presence of a dorso-ventral boundary and adaxial-abaxial interactions specifies leaf laminar margins (Byrne, 2012).

The margin tissue is notable in that it does not divide in the z-axis, maintaining the thallus edge as one cell in thickness, and does not diversify to produce any other cell types after its initial emergence out of the apical notch. We suggest a possible model where specification or not as margin cell is the first fate choice available to cells produced by the meristematic apical cells as they leave the apical notch. If the cell is at the edge, with one side not touching any other cells, then integration of signals would allow it to detect that its position must be at the thallus edge and to adopt a margin tissue fate. If, on the other hand, the cell is in contact with other cells on all sides, then it would not acquire margin cell fate and instead follows possible “inner” specification pathways, to produce other cell types such as air chambers, oil cells etc. An exception to this in normal development would be the formation of the second, distal set of margin tissue that emerges despite being in contact with other cells. After the second set of margin tissue emerges, no further new sets of margin tissues form (possibly due to inhibitory feedback effects) and the three-dimensional thalloid growth of the mature gemma proceeds, unless there are other influences such as elevated auxin levels (see below). In this way the margin tissue could play a role in defining the thalloid growth form.

The insertion site in line ET239-P64 was identified as being on chromosome 4, at site 24102312 in the MpTakv6.1 genome annotation (Figure 3 (m)). Potential candidate genes nearby are Mp4g21320 (Mapoly0090s0089, a cyclophilin-type peptidyl-prolyl cis-trans isomerase), Mp4g21310 (Mapoly0090s0090, a gene with no similarity to any other genes in the Genbank/NCBI database) and a third candidate gene Mp4g21300 (Mapoly0090s0091). Mp4g21300 is annotated in the *Marchantia polymorpha* v6.1 genome as being Mp1R-MYB17 and having significant sequence similarity to Myb transcription factors (highest similarity in Arabidopsis to Myb20 and Myb42). Mp1R-MYB17 has been noted to be strongly up-regulated in response to auxin (2,4-D) treatment from RNAseq experiments (Mutte *et al*., 2018; Flores-Sandoval *et al*., 2018). The activity of this gene has not been characterized in Marchantia, but Myb20 and Myb42 genes are known to be connected to lignification-related processes and bark formation in vascular plants (Geng *et al*., 2020; Wang and Dixon, 2012) and other Myb family genes are connected to cuticle formation in Marchantia (Albert *et al*., 2018). While bryophytes lack lignin, they do synthesize some lignin precursors in their cuticles (Bowman *et al*., 2017; Kolkas *et al*., 2022; Renault *et al*., 2017). This raises the possibility that the margin tissue cell wall composition may be different from the cell walls of the rest of the thallus. Interestingly, Mp4g21300 expression is significantly enriched in one unannotated cluster (Cluster 13) of the scRNA-seq analysis. Cluster 13 has Gene Ontology term enrichment for flavonoid and phenylpropanoid biosynthesis, as well as terms connected to cell wall biogenesis and polysaccharide biosynthesis. We speculate that Cluster 13 could be margin tissue. This would tally with Cluster 13 being a greater proportion of cells in older gemmae (31 days after germination), when the pattern of thallus growth would lead to relatively more margin tissue compared to younger, rounder gemmae (Wang *et al*., 2023). Combined with the auxin responsiveness of the trapped enhancer element (see below), and the possible functional role of margin tissues in Marchantia morphogenesis, this line provides a promising tool for future research.

### Margin Tissue Marker Response to Auxin Treatments

Application of exogenous auxin to Marchantia thalli induces aberrant growth to produce heavy ridges (Kato *et al*., 2017; Flores-Sandoval *et al*., 2015). To test whether the margin tissues are affected, ET239-P64 line gemma were grown on media containing 1µM NAA (Figure S3(a), (b)), 3µM NAA (Figure S3(c), (d)) or 5µM NAA (Figure S3 (e)-(g)). Elevated auxin levels resulted in the appearance of additional mVenus signal in a dose dependent manner. This additional signal is not restricted to the thallus edges but is found further in towards the central zone and thallus lobes compared to untreated control plants (Figure S3(h), (i)). Time lapse imaging revealed that the additional signal appears spontaneously within existing thallus cells, rather than being derived from production of extra fluorescent cells growing out from the apical notches (Movie S3). The cells displaying nuclear-localised mVenus do not necessarily have the characteristic “pillow-shaped” morphology of regular margin tissue cells, even at the thallus edge where they can possess irregular protrusions (Figure S3(g)).

In contrast, application of the auxin synthesis inhibitor yucasin resulted in a dramatic loss of margin tissue signal (Figure S3(j), (k)). This may be related to the wider effects of auxin synthesis inhibitors on the morphology of the cells at the apical notch (Flores-Sandoval *et al*., 2015). Gemmae grown on media supplemented with the auxin transport inhibitor TIBA did not show altered distribution of the margin tissue signal in the young gemmae (Figure S3 (l), (m)) but did exhibit a noticeable change in disrupting the formation of the additional set of margin tissue compared to untreated control plants (Figure S3 (n)-(s)). The z-axis split in the gemmae failed to occur and no air chambers were formed, even in gemmae grown for up to 2 weeks (e.g. Figure S3 (p)-(s)). This highlights how the formation of the second set of margin tissue is associated with the z-axis split and that this is a critical stage in producing the mature thallus morphology (Shimamura, 2016)

Validation experiments were performed to verify that approach taken in the auxin manipulation experiments did indeed affect gemmae growth and gene expression. Gemmae from line ET239-P54 were also grown in each treatment, showing that the auxin and inhibitor treatments did not alter enhancer trap signal non-specifically (Figure S4(a)). ET239-P54 marks oil cells, which are not known to be auxin-responsive. Further controls used a transgenic marker line with a fluorescent protein gene driven by the promoter of the auxin synthesis gene YUCCA2 (Figure S4(b)). MpYUC2 marker plants demonstrated predictable responses: reduced gene expression correlated to elevated auxin levels, and increased expression under inhibitor treatment (Figure S4(c)). It is noteworthy that in these plants the gene expression response was experienced uniformly across the gemma, rather than being limited to any one region or tissue type. This is consistent with previous qPCR analysis, showing that auxin treatment reduces MpYUC2 expression (Mutte *et al*., 2018).

The effects of manipulation of auxin levels on margin marker expression strongly suggests that the associated gene expression, and the formation of margin tissue, is auxin dependent. This is corroborated by the spatial localization of expression of the auxin transporter gene MpPIN1 (Figure S4(d)). Marchantia has only one copy of this canonical polar auxin transporter. In 0dpg gemma PIN expression is focused around the apical notch. As the gemma grows, there is scattered expression in the central zone, concentrated expression around the apical notch, and a chain of expression around the gemma edge. As the gemma grows further, the pattern of expression clearly shows MpPIN1 localized around the gemma edge, co-incident with the margin tissue region. PIN gene expression is known to be up-regulated by auxin (Tanaka *et al*., 2006; Paciorek *et al*., 2005). As with MpYUC2, the MpPIN1 marker line plants also displayed the predicted response, where elevated auxin treatment increased the distribution of MpPIN1 gene expression, particularly in the central zone of the gemma, correlated with NAA concentration (Figure S4(e)). In contrast, inhibitor treatment reduced MpPIN1 expression, most conspicuously around the apical notches of treated gemmae (Figure S4(e)). This is evidence for spatial heterogeneity of auxin in the gemma (Ishida *et al*., 2022).

Additionally, when over-expression of the gene MpLAXR (MpERF20) was induced, it was observed that cells around the gemmae edges divided strongly to form cellular masses (Ishida *et al*., 2022). It is plausible that MpLAXR (MpERF20) is acting as part of a regulatory system for margin tissue, with high auxin levels repressing expression to specify fate. These pieces of evidence point towards auxin being a signal in the specification of cells as “outer” (i.e. margin tissue) or “inner” (i.e. not margin) immediately after they are produced from the meristematic apical notch.

### Apical Notch/Meristem Marker lines

A further benefit of enhancer trap systems is to reveal finer scale anatomy of key, but poorly understood, features. The notch-like apical meristem of Marchantia provides a case in point. Thirteen lines were selected as having particularly strong, specific and reliable signal in and/or around the apical notch (Figure 4 and Figure S5). The growth and development of the notch signals were tracked from 0dpg until 3dpg for each line. The distribution of signal in each line was subtly different, in some cases being restricted to the immediate apical notch (e.g. ET239-P161 Figure 4(a), ET239-P156 Figure 4(b)), or being found at and around the apical notch (e.g. ET239-P49 Figure 4(c)), particularly along the flanks of the notch (e.g. ET239-P14 Figure 4(d)). In several cases signals were present throughout the gemma initially, but as gemma growth proceeded it became more concentrated at the apical notch before fading as cells aged (e.g. ET239-P133, Figure 4(e)). It should be noted that these lines could be interpreted as marking gene expression related to meristem-specific processes in the apical notch, or they may simply be markers of cell age, and therefore represent “proxies” of meristematic activity. A further range of markers for the apical notch region are demonstrated across the eight lines shown in Figure S5 (a)-(h).

**Figure 4.**
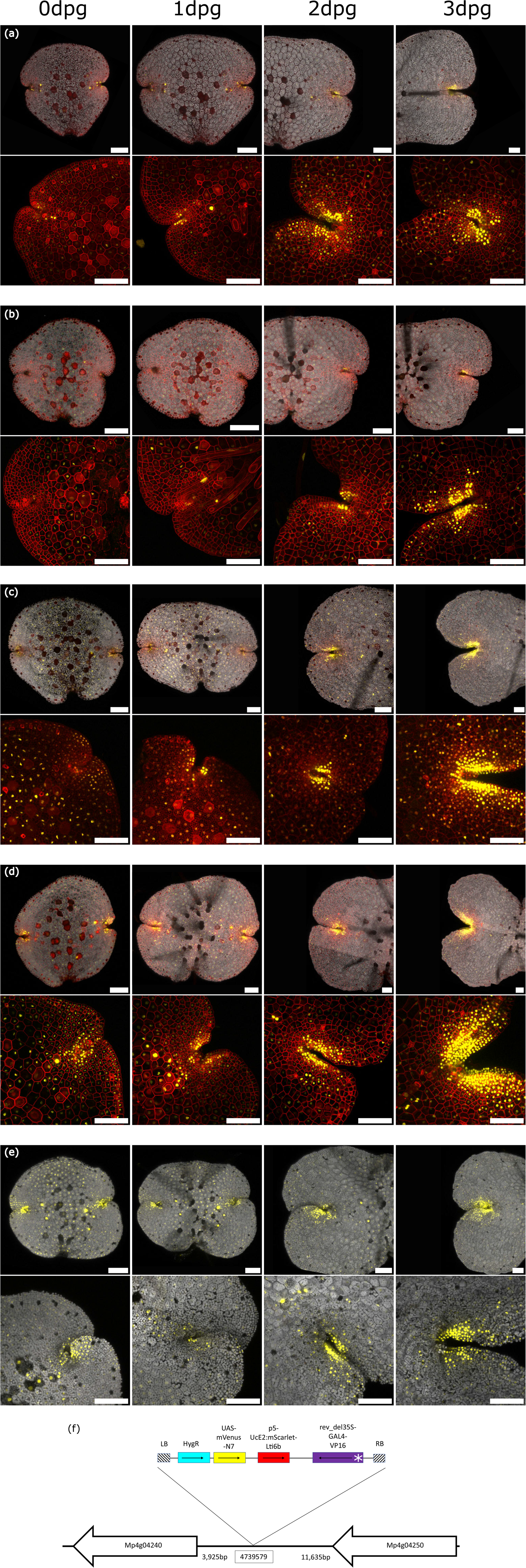
Selected marker lines for the apical notch region of the gemma. (a) ET239-P161, (b) ET239-P156, (d) ET239-P49, (d) ET239-P14, (e) ET239-P133. Top row in each sub-figure is the same gemma imaged at 0dpg, 1dpg, 2dpg and 3dpg. Bottom row in each sub-figure is higher magnification image of a different gemma from each line imaged at 0dpg, 1dpg, 2dpg, 3dpg. Scale bars = 100µm. Note that ET239-P133 (e) lacks the membrane signal, possibly due to disruption of the T-DNA insert in the genome. (f) Genomic location of the insertion site in an exemplar apical notch/meristem marker line, ET239-P161. The location of the insert site is on chromosome 4, nearby the putative candidate gene Mp4g04240 (Mapoly0044s0049), which is annotated as containing the embryophyte-specific domain of unknown function PTHR33782. Neighbouring genes are indicated as white arrows, starting at the gene’s start codon and pointing towards the gene’s stop codon. Schematics are not drawn to scale and distances are indicated in base pairs. The T-DNA map shows the relative positions of the minimal promoter (white asterisk), GAL4 enhancer (purple), mScarlet membrane marker (red), UAS-mVenus nuclear marker (yellow) and hygromycin resistance gene (cyan), as well as the left and right border sequences (hatched). Black arrows point from the start codon towards the stop codon for each gene.

The insertion site was identified in one line, ET239-P161 (Figure 4(a)), as an exemplar of an apical notch/meristem marker line. The insertion site was located within an intergenic region of chromosome 4 (site 4739579 in the MpTakv6.1 genome annotation). This is adjacent to the candidate gene Mp4g04240 (Mapoly0044s0049) (Figure 4(f)), which has been annotated in the *Marchantia polymorpha* v6.1 genome as having a domain of unknown function PTHR33782. Analysis of scRNA-seq data finds that expression of Mp4g04240 is enriched in the air pore cluster, although the authors do note that their analyses were limited by the fact that the proportion of cells sampled from near the notch was relatively low (Wang *et al*., 2023). While the molecular function of this gene and the PTHR33782 domain is unknown, BLAST searches against the EMBL/Genbank nucleotide and protein databases finds that it has sequence similarity to a set of genes present in all land plants, and that this gene family is unique to embryophytes. This putative candidate gene family therefore represents an intriguing candidate for further study in other plant groups and their meristems.

*Marchantia polymorpha* exhibits remarkable regenerative properties, and if both apical notch regions are excised from a gemma, one or more new apical notches will regenerate and emerge from the remaining tissue, usually within 96 hours (Binns and Maravolo, 1972; Kubota *et al*., 2013; Nishihama *et al*., 2015). This occurs after a burst of cell division, before one or more of these proliferative patches develops into a new meristematic region and recapitulates the full apical notch morphology (Nishihama *et al*., 2015). We examined the apical notch marker lines by cutting away the region surrounding the apical notches in each gemma by means of laser ablation (see Methods), and then monitored gemmae daily for changes in marker signal. In all cases the signals re-appeared, marked the regions of regenerating meristem and persisted in newly formed apical notches. Most (10 out of 13) lines had their first new marker signal appearing 24-48 hours after cutting (Figure 5(a)-(c) and Figure S6(a)-(d), (g), (i), (j), (l)). These results indicate that the lines mark processes that occur early in meristem formation/regeneration and in the acquisition of meristem fate. It is also apparent that not all patches of cell proliferation develop into fully active notches and maintain apical notch/meristem marker signals (arrows, Figure 5 (b), (c)). In these cases, the marker signal is observed to fade and disappear over time, consistent with the view that these enhancer trap lines are markers of active meristematic tissue.

**Figure 5.**
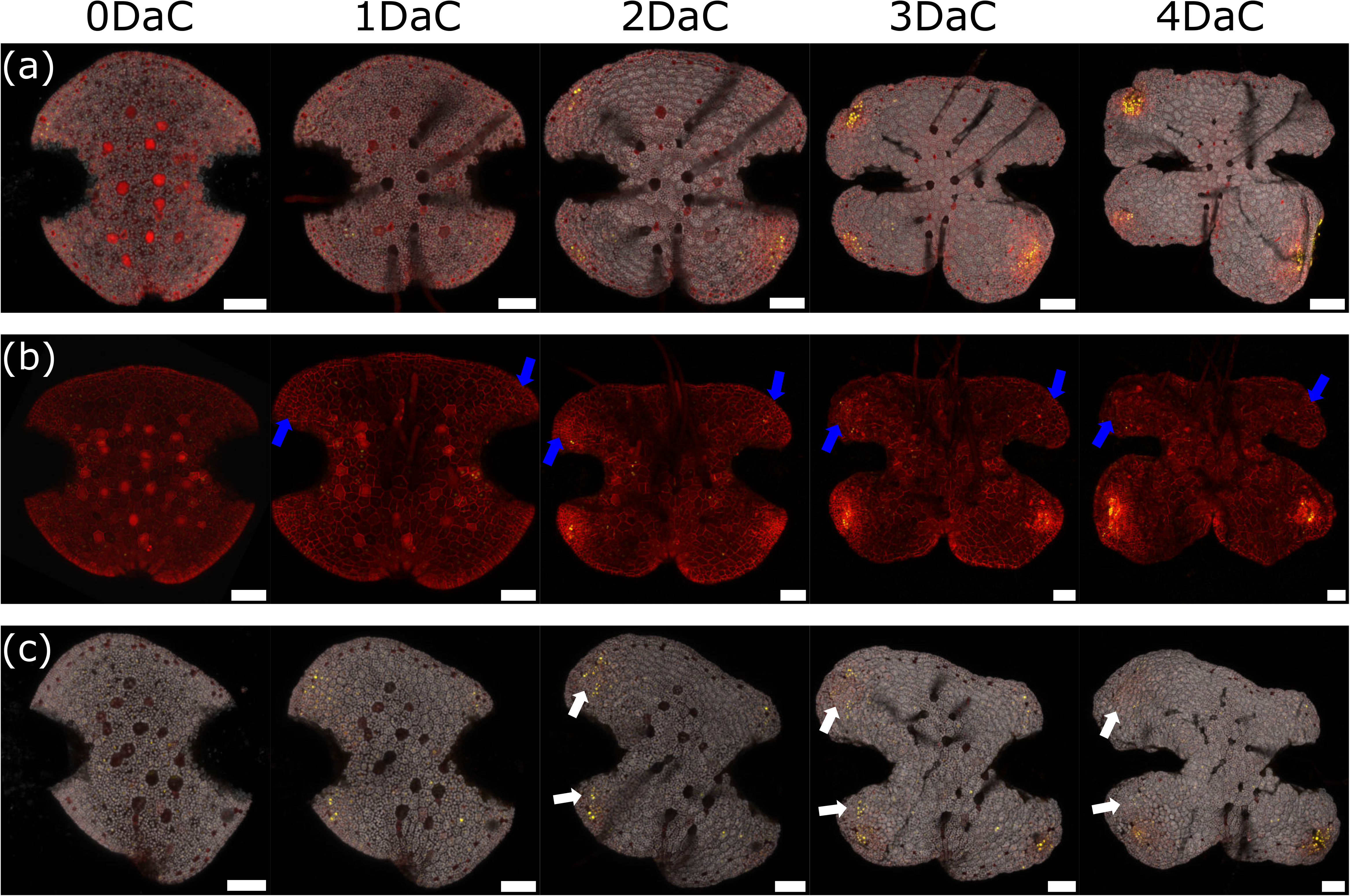
Timing of reappearance of apical notch/meristem marker signal after excision of the apical notches in selected lines. (a) Line ET239-P161, imaged 0-4DaC showing reappearance of nuclear mVenus signal after 24-48 hours that marks regions that regenerate full meristems by 4DaC. (b) Line ET239-P156 imaged 0-4DaC, showing reappearance of nuclear mVenus signal after 24-48 hours and full meristem re-emergence by 4DaC. Blue arrows show regions with cell divisions early on but where no marker signal emerges and persists, showing that marker signal is not simply related to cell division but instead is meristem-specific. The chlorophyll channel has been omitted for clarity. (c) Line ET239-P49 0-4DaC, showing reappearance of nuclear mVenus signal after 24-48 hours and full meristem re-emergence by 4DaC. White arrows show regions initially with cell division, but where cell division later stops and marker signal fade. This indicates that the dynamics of the marker relates specifically to meristem regeneration. Scale bars = 100µm.

The phytohormone auxin is a central player in plant meristem dynamics (Pernisová and Vernoux, 2021; Tanaka *et al*., 2006). Altering auxin levels in Marchantia by treatment with exogenous auxin or drugs that inhibit auxin synthesis is known to alter the apical notch region in particular (Maravolo and Voth, 1966). Therefore, we grew gemmae from each apical notch/meristem marker line supplemented with elevated auxin (1µM, 3µM or 5µM NAA) or with the auxin synthesis inhibitor yucasin (10µM) until 3dpg, after which the plants were imaged alongside untreated controls. The results are shown in Figure 6 and Figure S7. Each line displayed changes in spatial distribution of the nuclear signal under different auxin treatments. There were five broad classes of response: signal distribution changes relating to changes in notch architecture (e.g. ET239-P161, Figure 6(a); Figure S7(a)-(d)); reduced/no signal under both elevated auxin and inhibitor treatments (e.g. ET239-P156, Figure 6(b); Figure S7(e)-(h)); reduced signal under elevated auxin whereas inhibitor treatment only caused changes in spatial distribution (e.g. ET239-P49, Figure 6(c); Figure S7(i), (j)); elevated auxin response relating to notch architecture but inhibitor treatment eliminating/strongly reducing signal (e.g. ET239-P14, Figure 6(d); Figure S7 (k), (l)); large spatial changes in signal distribution under both elevated auxin and inhibitor treatment, unrelated to changes in notch architecture (e.g. ET239-P133, Figure 6(e); Figure S7(m)).

**Figure 6.**
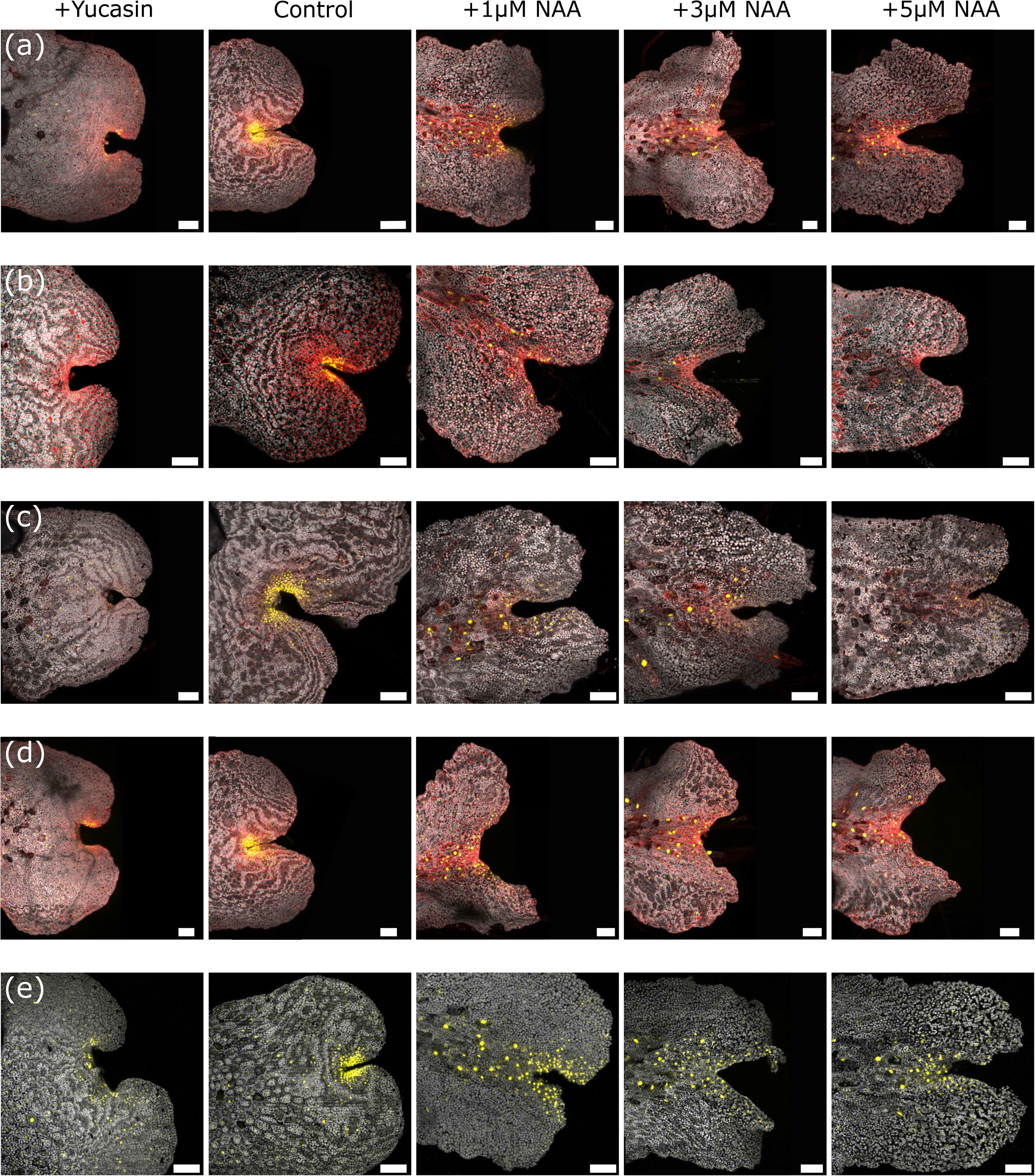
The effect of auxin manipulation of the growth media on apical notch/meristem marker line gemma in selected lines. (a) ET239-P161 showing signal distribution changes relating to changes in notch architecture. (b) ET239-P156 signal is reduced or eliminated under both elevated auxin and inhibitor treatments. (c) ET239-P49 shows reduced signal under elevated auxin whereas inhibitor treatment causes changes in spatial distribution. (d) ET239-P21 showing an elevated auxin response related to notch architecture but inhibitor treatment eliminates or strongly reduces signal. (e) ET239-P133 showing large spatial changes in signal distribution under both elevated auxin and inhibitor treatment, unrelated to changes in notch architecture. All gemma shown were imaged at 3dpg. From left to right +10µM yucasin, untreated control, +1µM NAA, +3µM NAA, +5µM NAA. Scale bars= 100µm.

It is notable that the four lines where alteration of the auxin environment markedly reduced nuclear mVenus signal (ET239-P75, ET239-P82, ET239-P127 and ET239-P156, see Figure 6(b) and Figure S7(e)-(h)) all show a pattern in the control plants of having marker signal expression close to the notch. This suggests that the enhancer elements trapped in these lines and the cells and tissues marked are involved in central, core meristem functions, where perturbations of auxin levels are extremely disruptive. These were also lines where marker signal took longest to re-appear after apical notch excision: ET239-P75 (3DaC), P82 (3DaC), P127(3-4DaC) and P156 (2DaC, same as in uncut control plants where signal appears at 2dpg) (see Figure 5(b) and Figure S6(e), (f), (h), (k)). This supports the view that the genes trapped in these lines are involved in central meristem processes that only occur in mature, fully-functioning apical notches. Further, the varied behaviour of meristem markers is evidence for proximodistal substructures in the Marchantia apical notch, which is likely more complex than previously recognized. The zonation may be more akin to the way angiosperm shoot apical meristems can be divided into peripheral zone, central zone and organizing centre (Pernisová and Vernoux, 2021), rather than a simpler model where the Marchantia meristem region is comprised of an apical cell surrounded by merophyte cells (Suzuki *et al*., 2020; Shimamura, 2016).

### Using Markers from Enhancer Trap Lines to Study Sporeling Development

Once marker lines had been characterized in the gemma, the same lines could be used to investigate tissues and cell types in other parts of the Marchantia life cycle. Spore germination and sporeling development provided a test case. While the series of morphological transitions in sporeling growth are broadly known (O’Hanlon, 1926), links with processes in gemmae growth and thallus regeneration (Nishihama *et al*., 2015) are unclear. While superficially similar structures are seen during development of sporelings, gemmae and mature thalli, there is scant knowledge of whether the underlying mechanisms are the same. Furthermore, it is poorly understood how important features in sporelings, such as the emergence of a thalloid growth form, are regulated (e.g. by phytohormones or photosynthesis) and how this relates to gemma growth regulation. Spores were produced by crossing wild-type plants with a marker line from each of the categories discussed above: rhizoids (ET239-P177), oil cells (ET239-P54), margin tissue (ET239-P64) and apical notch/meristem (ET239-P161). The spores were spread on agar plates (taken as the point of germination) and individual sporelings followed on subsequent days post-germination (dpg) in time course experiments to monitor the appearance and patterns of the nuclear mVenus signal. Here our unbiased markers provide ideal tools to interrogate and better understand sporeling development.

The earliest appearance of nuclear mVenus signal in sporelings from the notch/meristem marker line ET239-P161 (Figure 7(a)) was in protonema at 6dpg, with most plants showing patches of marker signal by 9dpg-10dpg. The appearance of the mVenus signal occurred after multiple cell divisions and the production of rhizoids in the early stages of spore germination (O’Hanlon, 1926). There was no apparent correlation between sporeling size and the timing of meristem marker appearance, nor was there any obvious morphological threshold to predict the timing or location of emergence. Protonema showed at least one patch of marker expression before proceeding to the prothallus stage, although in most cases multiple patches were present on the same protonema. Of these, one main patch proceeded to dominate and become the apical “pro-notch” of the main emerging prothallus, although in rarer cases two patches proceeded to become growth centres and multiple prothalli would grow from the same sporeling (Figure 7(a), yellow arrows). These results indicate that line ET239-P161 is not simply a marker of cell proliferation, but specifically marks meristematic tissue that can proceed to form the apical notch of a (pro)thallus.

**Figure 7.**
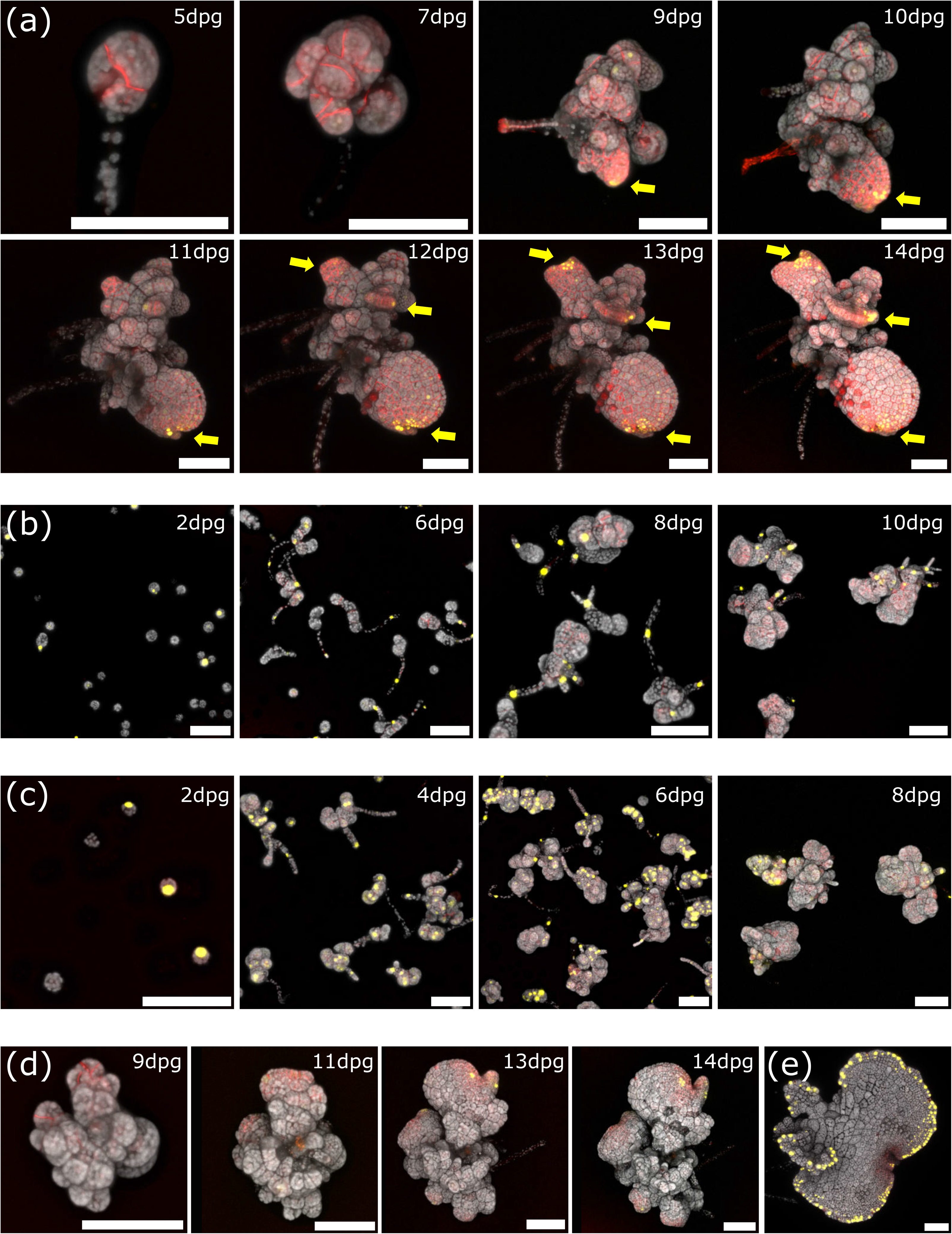
Sporeling development in enhancer trap lines. (a) Apical notch/meristem marker line ET239-P161 sporeling developmental sequence from 5dpg until 14dpg. Marker signal is not simply an indicator of cell division, but its appearance only precedes the prothallus stage and marks out the notch region of a new prothallus. Multiple “pro-notch” regions may be present in the same sporeling (yellow arrows). (b) Rhizoid marker line ET239-P177 sporelings. mVenus marker signal appears in the rhizoids and in those cells that form rhizoids. (c) Oil cell marker line ET239-P54 sporelings. Marker signal is present early in the germinating spores and throughout sporeling development, often in dense clusters and heterogeneously both within and between sporelings. (d) Margin tissue marker line ET239-P64 sporeling developmental sequence, imaged from 9dpg until 14dpg. The mVenus signal of the margin tissue only appears after the formation of an apical notch and the emergence of the prothallus stage. (e) ET239-P64 sporeling imaged at 12dpg displaying strong margin tissue marker signal around the prothallus edge. Scale bars= 100µm.

Further, the marker highlights a key step in sporeling development. There is a distinct difference between the protonema stage, which comprises callus-like photosynthetic tissue and rhizoids, and the prothallus stage, where differentiated meristematic tissue has become specified in a critical developmental transition. The emergence of an organised region of meristematic tissue is crucial for producing a thalloid morphology. The ability to mark and recognize the dynamic emergence of meristematic cells in sporelings is a major advance in understanding Marchantia development. This opens up new possibilities for using gemma and sporelings to better understand the molecular and cellular processes involved in meristem regulation in Marchantia (Hirakawa *et al*., 2019; Hirakawa *et al*., 2020), and plant meristems more generally (Schwartz *et al*., 2021; Kitagawa and Jackson, 2019).

A similar developmental sequence occurs during the regeneration of a full thallus from a small group of isolated non-meristematic cells (Figure S8). As with spore germination, the first set of divisions proceed slowly (Figure S8(a), (b)), and rhizoids emerge early on as the first new cell type (red arrow, Figure S8(c)). Other than the rhizoid cells, the photosynthetic portion of the regenerating plant proceeds to a morphologically undifferentiated callus stage marked by multiple cell divisions (Figure S8(d)). Out of this callus one or more patches of meristematic tissue emerge (as evidenced by the appearance of ET239-P161 marker signal, white arrows in Figure S8(e)-(i)). These meristematic patches proceed to produce a “protothallus” (Figure S8(j)), after which one or more meristematic regions grow to dominate and become the notches of a mature thallus. The regeneration of isolated thallus cells follows a scheme outlined for protoplasts (Ono *et al*., 1979; Bopp and Vicktor, 1988), however this does not require chemical methods to remove the cell wall, specialized liquid media for propagation, nor an additional time interval for *de novo* synthesis of a new cell wall. Additionally, isolated cells can be identified as being derived from a specific tissue type (in this case epidermis), and the deprogramming process can be directly followed. In contrast, regeneration from protoplasts includes the uncertain loss of cell properties during isolation and unknown origin of the progenitor cells. Using this single-cell regeneration technique, we found that cells from across all regions of a newly planted gemma, except the stalk scar, were equally capable of regenerating into new meristems. This is in contrast to regeneration studies on older thalli (10dpg-14dpg) which showed a bias towards regeneration from the apical edge of the ventral midrib epidermis (Nishihama *et al*., 2015; Ishida *et al*., 2022). This may be because gemmae initially lack dorsoventrality (Miller and Voth, 1962; Bowman *et al*., 2016), but it does suggest that cells throughout a gemma are capable of becoming adult stem cells, in contrast to, for example, Arabidopsis roots, where adult stem cells arise out the of the pericycle or pericycle-equivalent cells (Ikeuchi *et al*., 2016).

Germinating ET239-P177 spores showed marker expression in photosynthetic protonema cells before division to produce rhizoids, and in all differentiated rhizoids in the sporeling (Figure 7(b)). This indicates that sporeling rhizoids share some equivalence with those produced on both sides of early gemmae and ventral rhizoids of mature thalli. In contrast, observations of ET239-P54 spores during germination and sporeling development revealed broader patterns of marker expression (Figure 7(c). mVenus expression was observed in rhizoids and cells throughout the protonema and protothallus, arranged in irregular clusters throughout the sporeling. This suggests that the trapped regulatory element marked functions in wider biological processes occurring during sporeling development, not purely in oil cell formation.

Marker expression was observed in margin tissues during growth of ET239-P64 sporelings, consistent with observations that a distinct layer of peripheral cells is specified early in sporeling growth (O’Hanlon, 1926). No mVenus signal was observed in the early spore or protonema stages (Figure 7(d)). The first margin tissue marker signals were observed, after the emergence of recognisable meristematic regions (see below), generally 8dpg-12dpg. The morphology of the sporeling prothallus margin tissue cells is similar to those of the gemma, and distribution of marker expression in the margin tissue signal is the same, running around the periphery of the prothallus (Figure 7(e)). This indicates that specification of the margin tissue is closely associated with establishment of the thalloid form of growth.

Treatment of ET239-P64 spores with yucasin did not prevent the appearance of margin tissue marker expression in the prothallus, nor did it affect the distribution of signal versus control sporelings (Figure S9(a)-(d)), unlike in yucasin-treated gemma (Figure S3(h)-(k)). This indicates that the development of the prothallus and its margin tissue is different to some extent between germinating gemma and sporelings. Yucasin-treated sporelings in this case displayed greater levels of morphological development than in the more severe phenotypes observed in sporelings from knockout mutant lines for auxin synthesis or receptor genes (Eklund *et al*., 2015; Suzuki *et al*., 2023). This suggests that, for sporelings, the yucasin treatment incompletely inhibits auxin synthesis or that sporelings are capable of intracellular detoxification of yucasin. When ET239-P64 sporelings were grown under elevated auxin, most became stalled at the protonema stage, forming callus-like balls of cells lacking meristematic centres. No margin tissue signal emerged in these callus-type protonema (blue arrows, Figure S9(e)-(j)). Where the sporelings did proceed to the prothallus stage in elevated auxin treatments, margin tissue signal did appear. As in gemmae, prothalli grown under elevated auxin treatment showed additional margin marker signal in a broadly dose dependent manner. This demonstrates that expression of the marker for margin tissues is tightly linked to the emergence of the thalloid form, and that the induction of margin tissue marker signal by auxin occurs even at the sporeling stage (Figure S9(e)-(j)).

ET239-P161 spores were also grown under different treatments to manipulate auxin levels, with media supplemented with NAA, the auxin synthesis inhibitor yucasin or the auxin transport inhibitor TIBA. This was to examine how changes in auxin levels influence meristem emergence and in particular the transition from generalized cell division in the protonema to meristem-localised cell division in the prothallus. Compared to untreated control sporelings (Figure 8(a)-(c), yucasin treatment initially resulted in the formation of enlarged cells after spore germination (Figure 8(d), (e)), slowing the early development of the protonema. However, yucasin treatment did not prevent or slow the emergence of meristematic regions (as designated by the appearance of the marker signal, Figure 8(f)). Later development to the prothallus stage was not affected, and mature sporelings were comparable to the control plants (Figure 8(p)). As noted for ET239-P64 sporelings, these phenotypes are milder than is seen in knockout mutants for auxin synthesis or receptor genes (Eklund *et al*., 2015; Suzuki *et al*., 2023). TIBA treatment resulted in the appearance of elongated cells during early germination, but with less cell division so that sporelings at the protonema stage are generally smaller compared to control plants (Figure 8(g), (h)). The cell morphology was reminiscent of yucasin-treated protonema cells (c.f. Figure 8(e) and (h)), albeit markedly smaller. The smaller size of the protonema did not affect the emergence of meristem signal nor the transition to the prothallus stage (Figure 8(i)), further supporting the finding that this developmental transition was not dependent on protonema size or morphology. The normal developmental progress of TIBA-treated sporelings is in contrast to *Mppin1* knockout mutant sporelings (Fisher *et al*., 2023), where a significant proportion became stalled at the protonema stage (termed ‘blob’ stage by Fisher and colleagues). This could indicate incomplete inhibition of auxin transport in sporelings, a sporeling-specific TIBA detoxification pathway, or that MpPIN1 knockout affects sporeling development more widely than in disrupting auxin transport alone. By 20dpg, TIBA-treated sporelings were overall smaller than control plants, but had similar morphology (Figure 8(p)). That inhibition of auxin synthesis and transport produces elongated or enlarged cells at the protonema stage is similar to the effects of growing sporelings under a phytochrome-inactive Red/Far-Red light cycle (Nishihama *et al*., 2015). This is reminiscent of the overlap between auxin and shade responses in other plant species (Iglesias *et al*., 2018).

**Figure 8.**
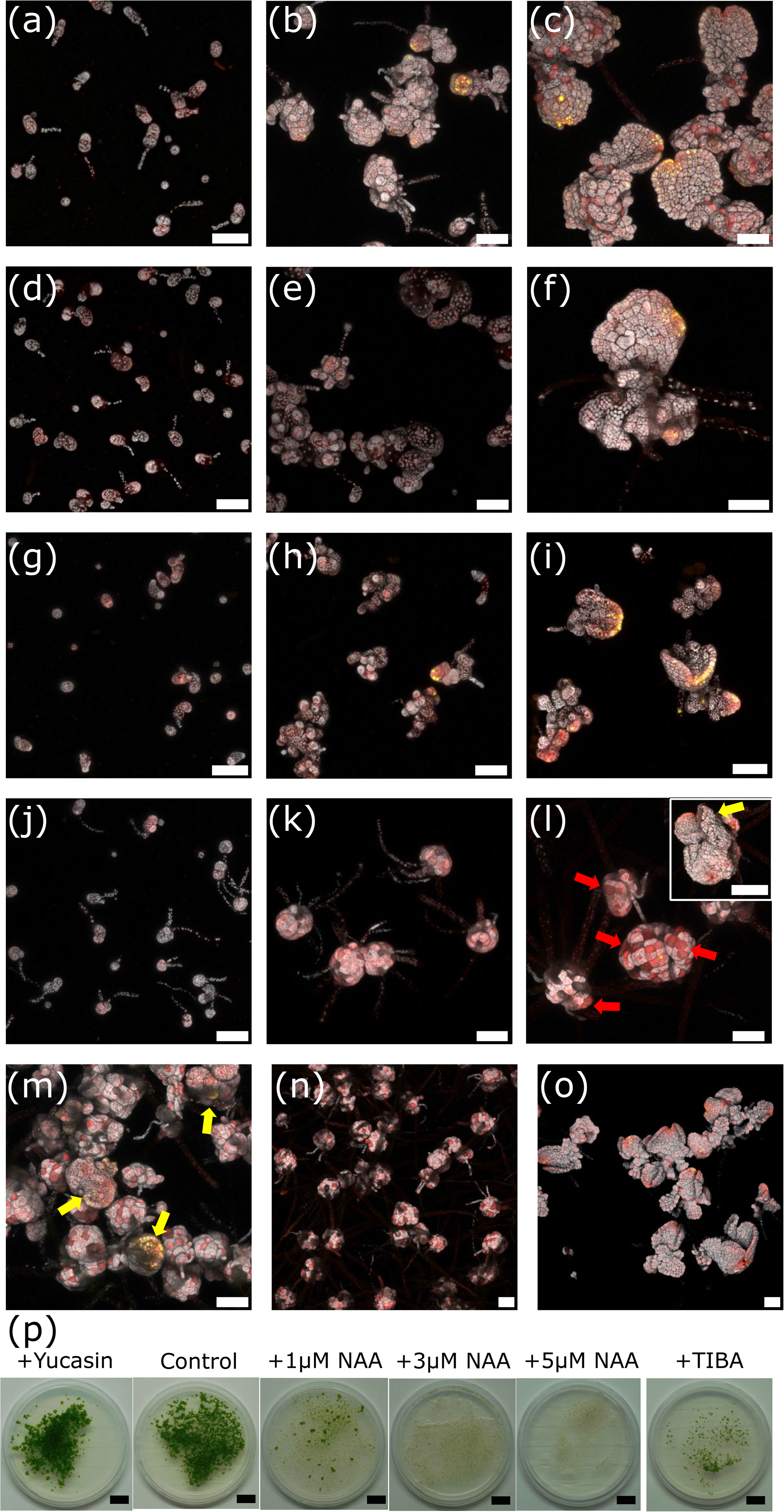
Effect of auxin manipulation on sporeling germination and development in apical notch/meristem marker plants. Control sporelings grown on untreated media imaged at (a) 4dpg, (b) 8dpg, (c) 12dpg. mVenus signal marks meristematic regions. Sporelings grown on +10µM yucasin media imaged at (d) 4dpg, (e) 8dpg, (f) 12dpg. Immediately after germination and in early protonema development yucasin-treated sporelings have enlarged vacuolated cells, however later protonema development and prothallus formation proceeds as normal, with emergence of a meristematic “pro-notch”. Sporelings grown on +100µM TIBA media imaged at (g) 4dpg, (h) 8dpg, (i) 12dpg. Initially sporelings have somewhat vacuolated cells and overall sporelings are slightly smaller, but there is no effect on prothallus formation and meristem emergence. Sporelings grown on +3µM NAA media imaged at (j) 4dpg, (k) 8dpg, (l) 12dpg. Early on in development, sporelings grown under elevated auxin treatment produce more rhizoids. Development then becomes stalled at the protonema stage, forming large callus-like masses of cells that contain rhizoids, photosynthetic cells and cells containing membranous organelles (red arrows). Only occasionally do sporelings proceed to the prothallus stage, with sporelings only producing one or at most two meristem regions (inset, yellow arrow). Sporelings imaged at 12dpg that were grown on +1µM NAA (m), +5µM NAA (n) or Control media (o). The numbers of sporelings that proceed to the prothallus stage versus those stalled at the protonema stage appears to be inversely related to the auxin concentration of the media. Where auxin-treated sporelings progress to the prothallus stage, only one or at most two meristem regions appear per prothallus (yellow arrows (n)). (p) Sporelings imaged at 20dpg that were grown on: +10µM yucasin, control, +1µM NAA, +3µM NAA, +5µM NAA, and +100µM TIBA media. Yucasin treatment produces no qualitative difference from untreated controls. TIBA treatment produces smaller sporelings. In contrast, even by 20dpg most auxin treated sporelings are dead or remain at the protonema stage, with this effect greater at higher auxin concentrations. All sporelings shown were grown from ET239-P161 spores. White scale bars= 100µm. Black scale bars= 1cm.

Auxin-treated spores showed increased rhizoid growth during the early stages of development (Figure 8 (j)). Protonema growth continued unaffected during the early stages, however development became stalled at this stage. The result was the production of large, ball-shaped protonema consisting of callus-like cells (Figure 8(k)), which (other than rhizoids or rhizoid precursor cells) appeared to be a poorly differentiated mass of dividing photosynthetic and cells with prominent membranous organelles (red arrows, Figure 8(l)). Escape from this stage was possible (e.g. inset Figure 8(l)) but depended on the exogenous auxin concentration. The appearance of meristem marker signal and the transition to the prothallus stage observed to be more frequent at 1µM NAA (Figure 8(m)) versus being rare (but present) at 5µM NAA (Figure 8(n)), compared to untreated control sporelings of the same age (Figure 8(o)). Once again, there was no link between protonema size and meristem emergence, nor were there any obvious morphological predictors for those plants that would make the transition to prothallus stage, until than the appearance of the meristem marker signal. The ‘stalling’ effect auxin has on sporeling development is still apparent at 20dpg (Figure 8(p)), with sporelings at higher auxin concentrations unable to complete their development and eventually dying. The effect of auxin manipulation is in contrast to the meristematic regeneration process in gemma, where treatment with auxin synthesis or transport inhibitors causes widespread cell division but prevents the emergence of full meristems, and the application of exogenous auxin inhibits any proliferative cell division at all in cut thallus pieces (Maravolo and Voth, 1966; Binns and Maravolo, 1972; Ishida *et al*., 2022). Therefore, while the development of thalli from both spores and isolated cells follows the same broad set of stages, the underlying processes differ in their response to auxin manipulation treatments (Flores-Sandoval *et al*., 2015; Eklund *et al*., 2018; Eklund *et al*., 2015; Ishida *et al*., 2022; Suzuki *et al*., 2023; Fisher *et al*., 2023). Superficial morphological similarities but underlying mechanistic differences between regenerating thalli and sporelings have also been noted for responses to Red/Far-Red light and sucrose supplementation (Nishihama *et al*., 2015). Discovering and understanding further developmental similarities and differences between these two life stages represents a promising avenue for future research.

## Conclusions

The creation of enhancer trap lines for *Marchantia polymorpha* has generated useful resources for use by the Marchantia research community. As well as the anatomical features of early gemma and spore development outlined above, the enhancer trap library contains resources for studying other tissue types, such as air pores and chambers, specific rhizoid types or the architecture of the gemma central zone. Furthermore, as the Marchantia genome has a moderate GC content of ∼45% (Marks *et al*., 2019; Bowman *et al*., 2017), the distribution of insertion sites should theoretically be random (Shima *et al*., 2016). The current screen was far from saturating and could be easily expanded by adoption of high throughput transformation and screening methods. This would likely reveal additional lines with novel cell markers. Targeted screening (i.e., searching for cell markers expressed specifically at certain developmental stages or growth conditions) could be repeated and applied to investigating other aspects of Marchantia development and gene expression, for example in sexual reproduction, release of gemma dormancy or responses to abiotic stress. This could be used to probe new cell interactions and for gene identification in tissues of interest. Further, the enhancer trap scheme employs the GAL4-VP16 gene as the primary target. This synthetic transcription factor has no natural counterpart in plant cells, and acts to amplify expression of the fluorescent marker gene. GAL4-VP16 can be used to drive ectopic expression of any other chosen gene fused to a suitable GAL4 promoter, and the pattern of expression will mirror expression of the marker gene. Similar approaches arising from a root-focused *Arabidopsis* enhancer trap project (Haseloff, 1999) have provided new tools and discoveries about other aspects of plant development (Laplaze *et al*., 2005; Wenzel *et al*., 2012; Radoeva *et al*., 2016).

The enhancer trap lines we have generated have allowed a better definition of fundamental processes in Marchantia, such as the stages of spore development. The appearance of meristematic regions was revealed to be distinct from simple localised cell division - a finding facilitated by the use of an unbiased collection of meristem markers such as those generated here. We have also shown that in the spore germination process key transitions are inhibited by applying exogenous auxin, but with smaller effects from pharmacological inhibition of auxin synthesis or transport. This contrasts with gemma regeneration and notch morphology which are heavily influenced by elevated auxin, or perturbations to auxin synthesis or transport. The enhancer trap lines will allow finer-scale studies of notch architecture, signalling and regeneration in Marchantia, and will complement other approaches that rely on the analysis of plants at the resolution of individual cells and tissues, including single-cell transcriptomics, promoter libraries and mutant phenotype analysis. This in turn will provide valuable insights and tools for plant science as a whole, owing to the conservation of transcription factors and genetic pathways throughout the embryophytes (Yamaoka *et al*., 2018; Bowman *et al*., 2017; Lu *et al*., 2020; Eklund *et al*., 2018)

## Supporting information

Legend for Movies S1, S2 and S3

## Acknowledgements

We would like to thank Satoshi Naramoto (Hokkaido University, Japan) for kindly providing the 5’ promoter sequence for MpYUC2 to create the vector construct. We thank members of the lab working with Marchantia, particularly Bernardo Pollak, Elizabeth Forsythe, Jenna Rever, Harriet Kempson, Sze Wai Tse and Eftychios Frangedakis, for their advice and support. This work was funded as part of the BBSRC/EPSRC OpenPlant Synthetic Biology Research Centre Grant BB/L014130/1 to J.H., BBSRC BB/F011458/1 for confocal microscopy to J.H. and BBSRC Research Studentship for M.R. (1943399).

---

**Figure S1.**
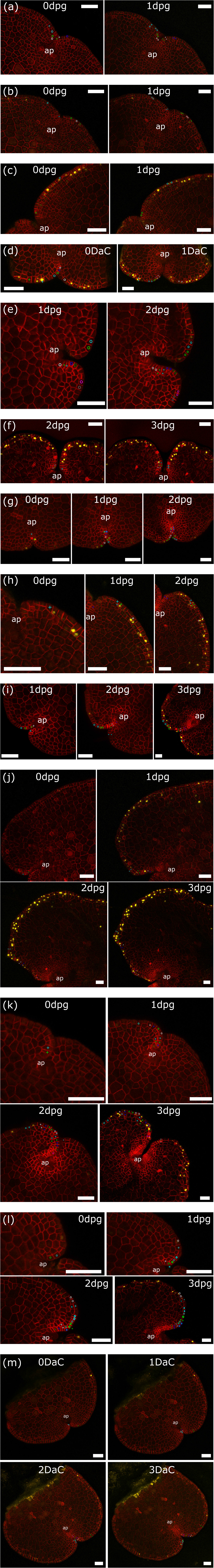
Tracking cell division in the margin tissue cells located at the edge of the gemma. The margin tissue cells located right at the gemma edge were monitored for division during a time course with daily confocal imaging. Cell division events were identified by the formation of new cell membranes, as shown by the mScarlet marker, since the previous image. The tracking finds that these cells can divide both parallel to the gemma edge (i.e. transverse, denoted by plus, +) and perpendicular to the gemma edge (i.e. anticlinal, denoted by circle, o), but not in the z axis (i.e. longitudinally). Cells can undergo transverse division after anticlinal division (denoted by asterisk, *) and vice versa (denoted by circled plus ⊕), or repeating anticlinal division (bold circles **o**). Most cell division in the margin tissue occurs around the apical notch (ap), with margin tissue cells further away from the notch being mitotically inactive. Cells are marked with a symbol when they have divided in the next time frame, with mother and daughter cells sharing the same colour symbols (note asterisks, denoting anticlinal followed by transverse division, are put outside the mother cell for clarity). (a) imaged 0dpg (left) and 1dpg (right). (b) imaged 0dpg (left) and 1dpg (right). (c) imaged 0dpg (left) and 1dpg (right). (d) imaged 0DaC (left) and 1DaC (cutting done on 0dpg gemma). (e) imaged 1dpg (left) and 2dpg (right). (f) imaged 2dpg (left) and 3dpg (right). (g) imaged 0-2dpg. (h) imaged 0-2dpg. (i) imaged 1-3dpg. (j), (k) and (l) imaged 0-3dpg. (m) imaged 0-3 DaC (cutting done on 0dpg gemma). All gemma shown are from the margin tissue marker line ET239-P64. The chlorophyll autofluorescence channel has been omitted for clarity in all images. Scale bars= 50µm.

**Figure S2.**
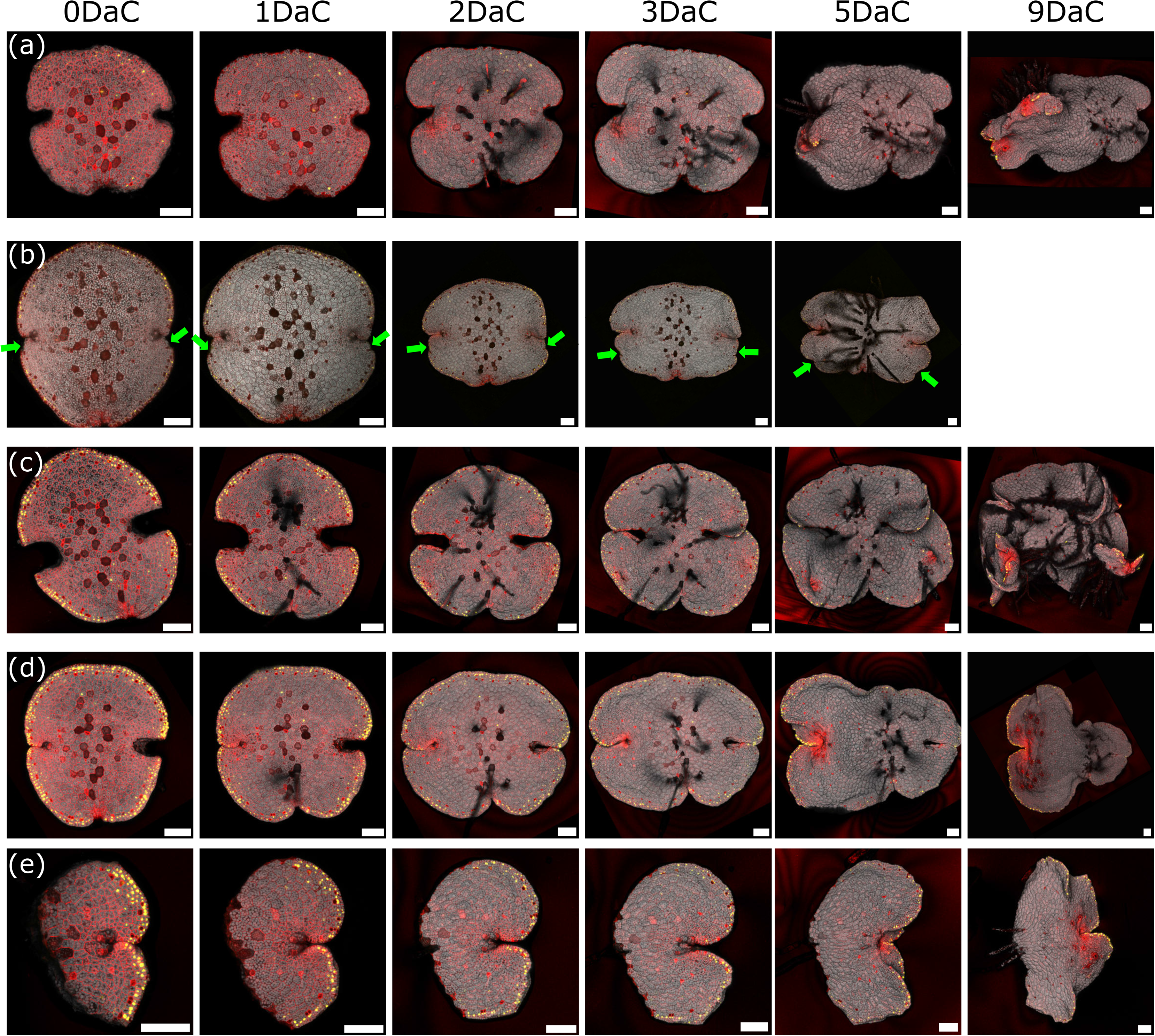
Laser ablation cutting and ventral view demonstrates that line ET239-P64 marks the cells that make up the margin tissue. (a) Gemma with all edges removed. (b) Gemma with small portion of edge removed (marked by green arrows) (c) Gemma with both notches removed but all edges intact (e) Gemma with one notch removed, other notch and all edges intact (e) Isolated gemma notch. Scale bars= 100µm.

**Figure S3.**
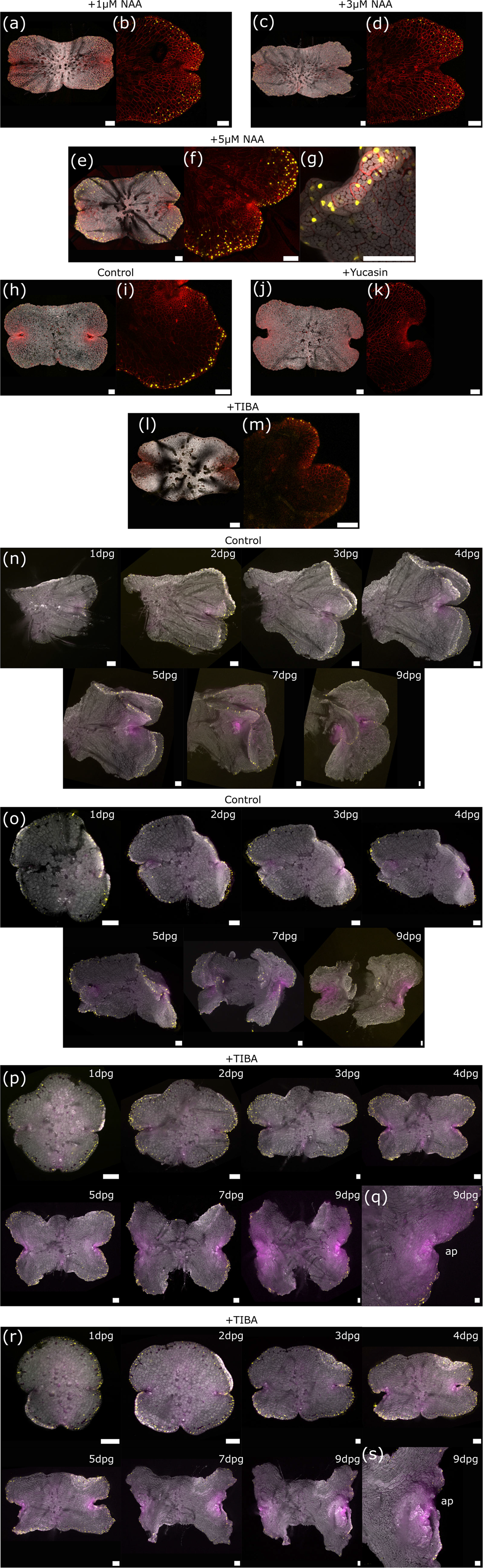
The effect of auxin manipulation of growth media on the margin tissue marker in line ET239-P64. (a), (b) +1µM NAA treated gemmae at 3dpg. (c), (d) +3µM NAA treated gemmae at 3dpg. (e), (f), (g) +5µM NAA treated gemmae at 3dpg. (g) is a close up image of a +5µM NAA treated plant showing cells exhibiting marker signal deeper inside the thallus and with irregular protrusions. (h), (i) Untreated control gemmae at 3dpg. (j), (k) +10µM yucasin treated gemmae at 3dpg. (l), (m) +100µM TIBA treated gemmae at 6dpg. Time course of gemma either untreated control ((n), (o)) or treated with 100µM TIBA ((p), (q), (r), (s)) imaged by Leica M205 FA stereomicroscope. mVenus channel is shown in yellow, mScarlet channel in magenta and chlorophyll autofluorescence chanel in grey. From left to right: 1dpg, 2dpg, 3dpg, 4dpg, 5dpg, 7dpg, 9dpg, 9dpg (close up of notch of 9dpg plant shown in (q) and (s), ap marking the apical notch). The first row of margin tissue is intact in both conditions, as shown by the marker signal being present. However, there is no formation of the second row of margin tissue, no z-axis split and no air chambers form even 7 days after removal from the gemma cup, in comparison to the normal development of the untreated control plants. Scale bars= 100µm.

**Figure S4.**
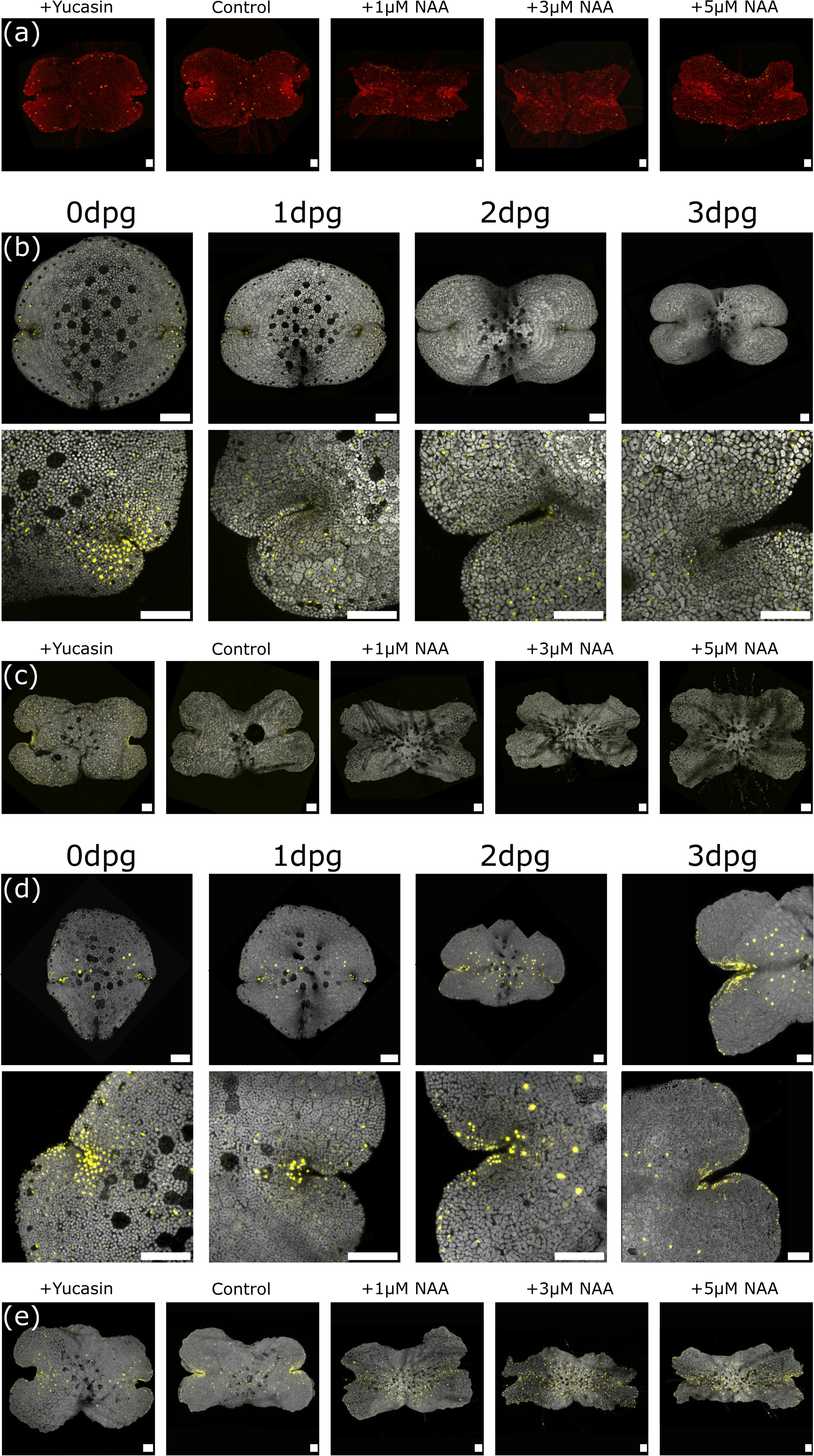
Effect of auxin manipulation on ET239-P54 (oil cell), MpYUC2 and MpPIN1 marker lines. (a) ET239-P54 marker line gemmae at 3dpg grown on media supplemented with +10μM yucasin, untreated control, +1μM NAA, +3μM NAA or +5μM NAA. Auxin manipulation does not appreciably change oil cell formation nor marker signal expression. (b) Characterization of MpYUC2 marker line. The top row is the same gemma imaged at 0dpg, 1dpg, 2dpg, 3dpg. mTurquoise channel shown in yellow. The bottom row is a higher magnification image of the area around the apical notch of different gemmae at 0dpg, 1dpg, 2dpg, 3dpg. mTurquoise channel shown in yellow. (c) MpYUC2 marker line gemmae at 3dpg grown on media supplemented with +10μM yucasin, untreated control, +1μM NAA, +3μM NAA or +5μM NAA. Inhibiting auxin synthesis strongly up-regulates MpYUC2 expression, whereas application of exogenous auxin decreases MpYUC2 expression. The effects are observed all across the gemma. mTurquoise channel shown in yellow. (d) Characterization of MpPIN1 marker line. The top row is the same gemma imaged at 0dpg, 1dpg, 2dpg, 3dpg. The bottom row is a higher magnification image of the area around the apical notch of different gemmae at 0dpg, 1dpg, 2dpg, 3dpg. These images show the prominent MpPIN1 expression in the margin tissue region, particularly illustrating the formation of the second row of margin tissue at 3dpg. (e) MpPIN1 marker line gemmae at 3dpg grown on media supplemented with +10μM yucassin, untreated control, +1μM NAA, +3μM NAA or +5μM NAA. Inhibiting auxin synthesis causes lower MpPIN1 expression, whereas application of exogenous auxin strongly increases MpPIN1 expression, particularly in the central zone of the gemmae. Scale bars =100μm.

**Figure S5.**
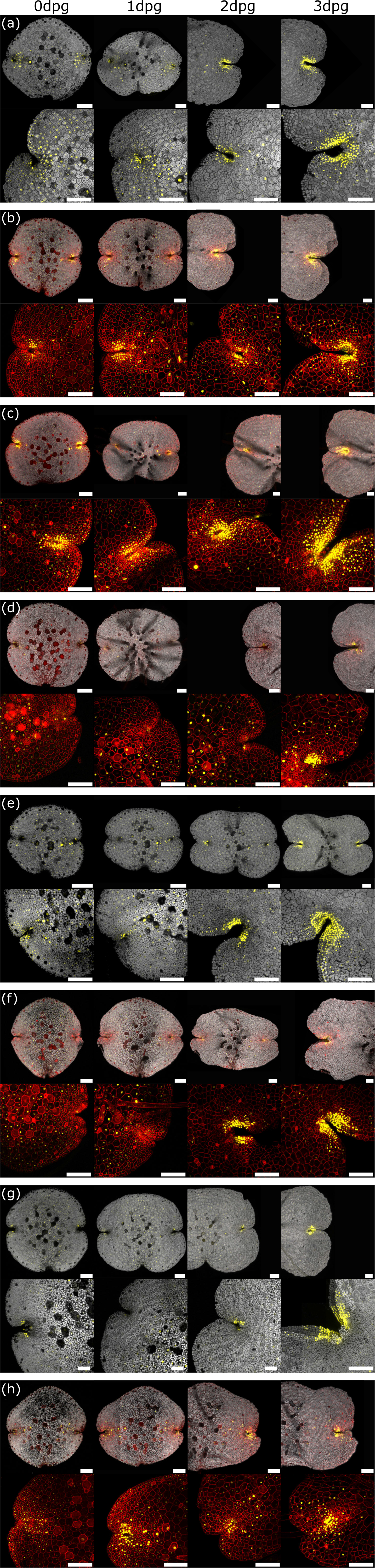
Apical notch/meristem marker lines. (a) ET238-P25 (b) ET239-P21, (c) ET239-P33, (d) ET239-P75, (e) ET239-P82, (f) ET239-P125, (g) ET239-P127, (h) ET239-P153. Top row in each sub-figure is the same gemma, imaged at 0dpg, 1dpg, 2dpg, 3dpg. Bottom row in each sub-figure is higher magnification image of different gemmae from each line, imaged at 0dpg,1dpg, 2dpg, 3dpg. Scale bars= 100µm.

**Figure S6.**
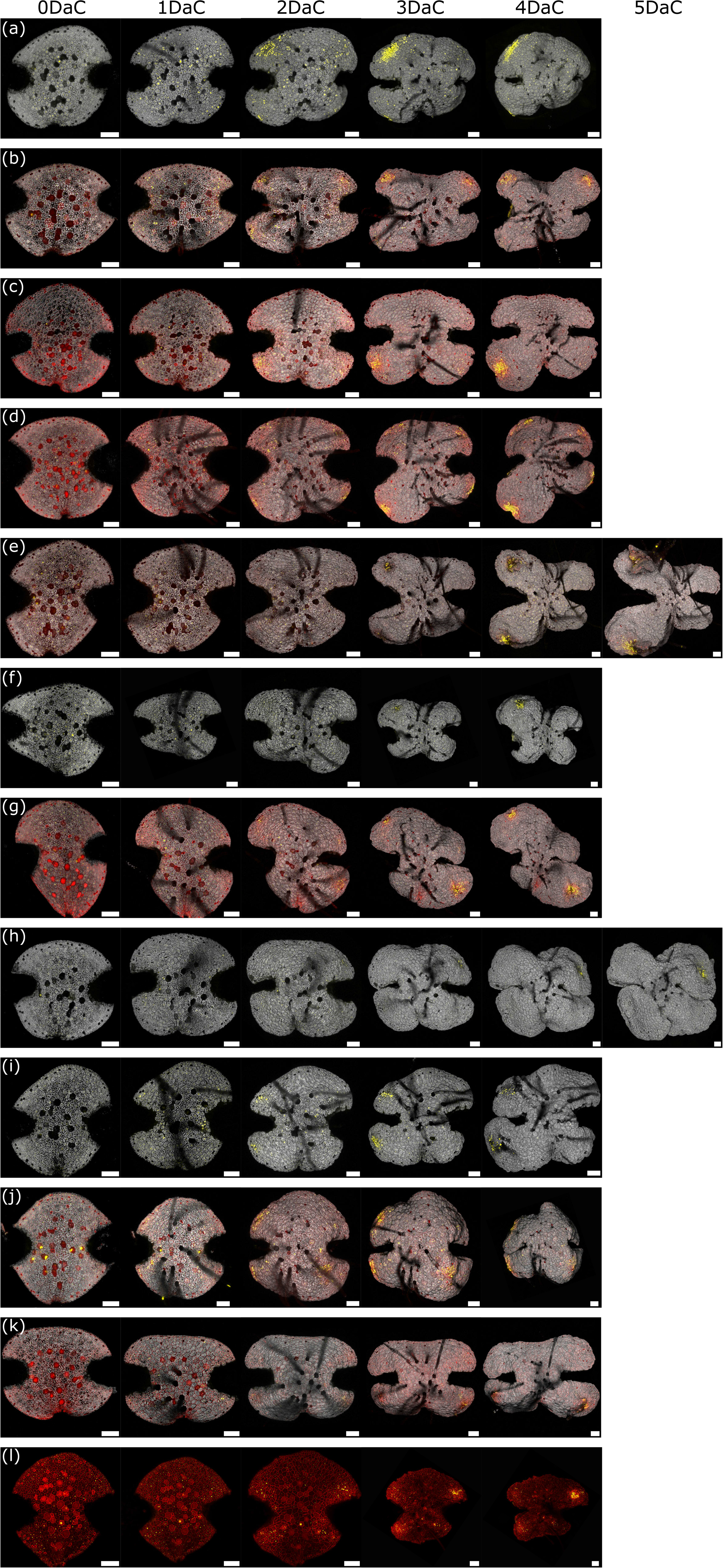
Timing of the reappearance of apical notch/meristem marker signal after excision of the apical notches. (a) ET238-P25 imaged from 0DaC until 4DaC. (b) ET239-P14 imaged from 0DaC until 4DaC. (c) ET239-P21 imaged from 0DaC until 4DaC. (d) ET239-P33 imaged from 0DaC until 4DaC. (e) ET239-P75 imaged from 0DaC until 5DaC. (f) ET239-P82 imaged from 0DaC until 4DaC. (g) ET239-P125 imaged from 0DaC until 4DaC. (h) ET239-P127 imaged from 0DaC until 5DaC. (i) ET239-P133 imaged from 0DaC until 4DaC. (j) ET239-P153 imaged from 0DaC until 4DaC. (k) ET239-P156 imaged from 0DaC until 4DaC (chlorophyll channel included, cf. Figure 5(b)) (l) ET239-P161 imaged from 0DaC until 4DaC (chlorophyll channel omitted, cf. Figure 5 (a)). Scale bars= 100µm.

**Figure S7.**
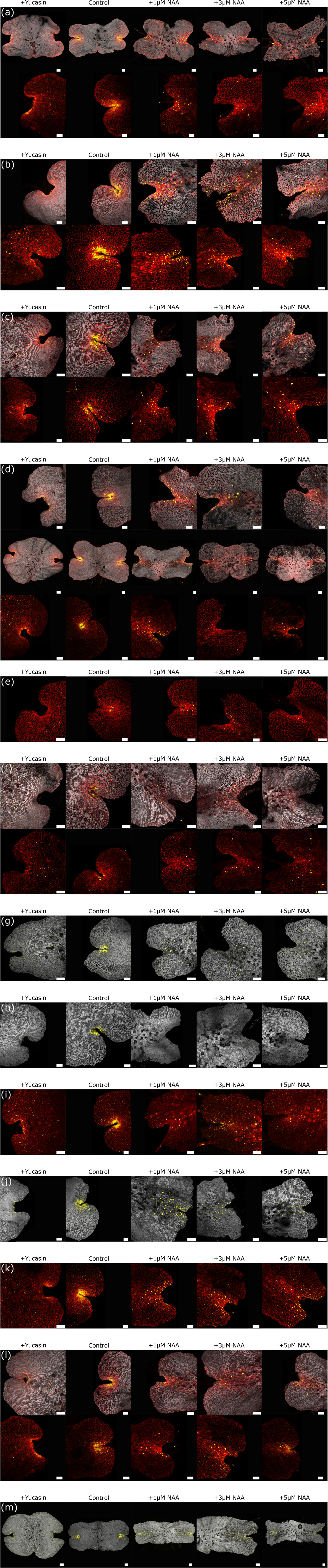
Auxin manipulation treatments of apical notch/meristem marker line gemmae. (a) ET239-P161 whole gemma images in top row, with chlorophyll channel omitted in images of a different gemma in bottom row. (b) ET239-P33; with chlorophyll channel omitted in images of a different gemma in bottom row. (c) ET239-P153; with chlorophyll channel omitted in images of a different gemma in bottom row. (d) ET239-P125; whole gemma images of a different gemma in middle row; chlorophyll omitted images of a different gemma in bottom row. The lines shown in (a)-(d) show signal distribution changes in respone to auxin manipulation that relates to changes in notch architecture. (e) ET239-P156 with chlorophyll channel omitted. (f) ET239-P75; with chlorophyll channel omitted in images of a different gemma in bottom row. (g) ET239-P82. (h) ET239-P127. The lines shown in (e)-(h) have signal that is dramatically reduced or eliminated under elevated auxin treatments and auxin synthesis inhibitor treatment. (i) ET239-P49 with chlorophyll channel omitted. (j) ET238-P25. The lines shown in (i) and (j) have reduced signal under elevated auxin treatments, whereas auxin synthesis inhibitor treatment causes changes in spatial distribution of signals. (k) ET239-P21 with chlorophyll channel omitted. (l) ET239-P14; with chlorophyll channel omitted in images of a different gemma in bottom row. The lines shown in (k) and (l) has a response to elevated auxin treatments that relates to notch architecture, but auxin synthesis inhibitor treatment eliminates or markedly reduces signal. (m) ET239-P133 whole gemma images. This line shows large spatial changes in signal distribution under both elevated auxin and inhibitor treatment, that appears to be unrelated to changes in notch architecture. All gemma shown were imaged at 3dpg. In all sub-figures from left to right: +10µM yucasin, untreated control, +1µM NAA, +3µM NAA, +5µM NAA. Scale bars= 100µm.

**Figure S8.**
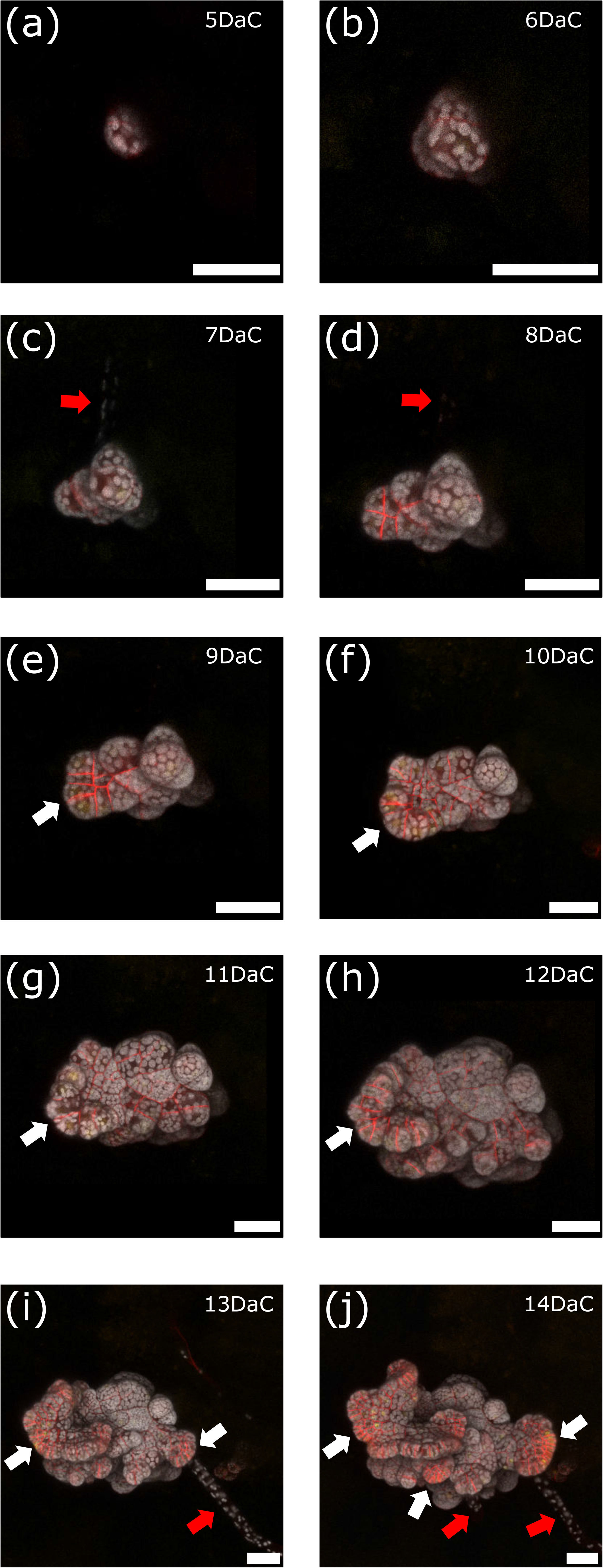
Developmental sequence of thallus regeneration from a small cluster of isolated gemma cells. Cells were isolated from the periphery of a 0DaC gemma from the apical notch/meristem marker line ET239-P161 by laser ablation. Images of the cells shown were taken daily from 5DaC (a) until 14DaC (j). Initially cell division proceeds with no marker signal apparent, to form a callus-like mass of photosynthetic cells with occasional rhizoids (red arrows). Marker signal appearance precedes and marks out the regenerating meristem region (white arrows) that will go on to form the apical notch of a new thallus. Scale bars= 50µm.

**Figure S9.**
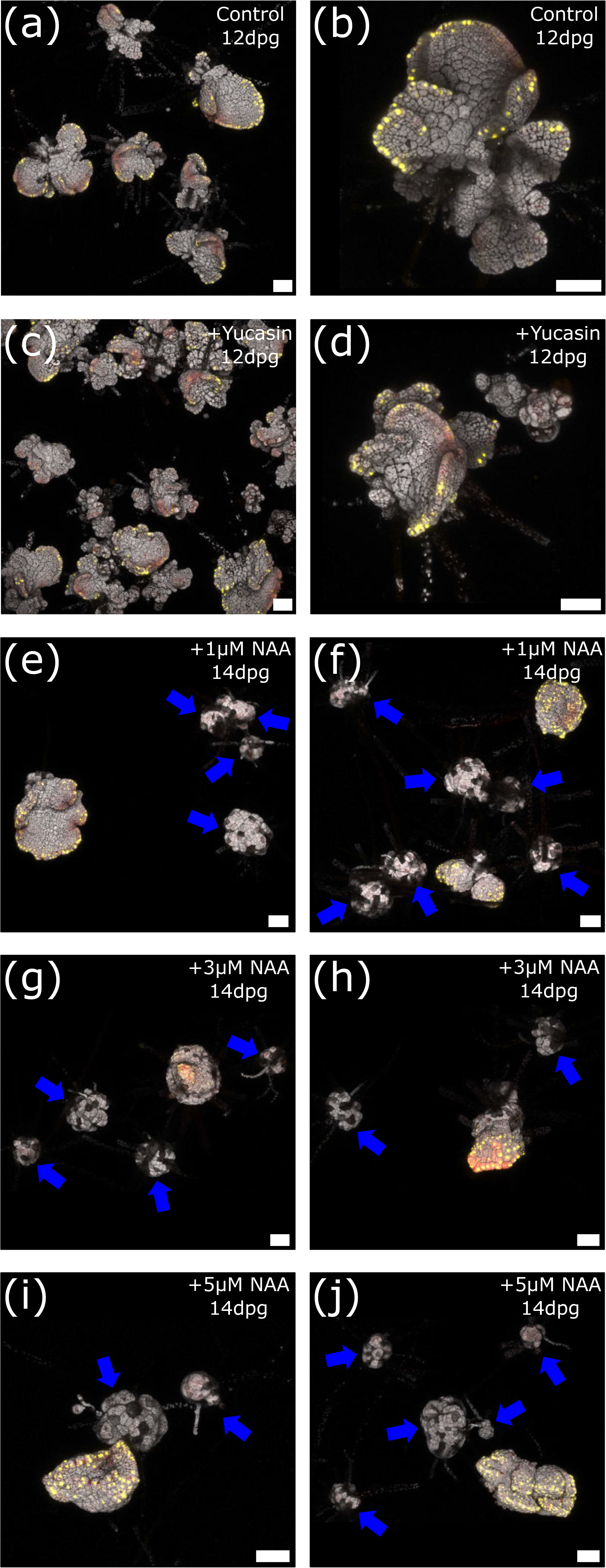
Effect of auxin manipulation on spore germination and development in margin tissue marker (ET239-P64) sporelings. (a), (b) Sporelings imaged 12dpg at the prothallus stage grown on control media. (c), (d) Sporelings imaged 12dpg at the prothallus stage grown on +10µM yucasin media. These show normal appearance of the prothallus and margin tissue marker signal. Sporelings imaged at 14dpg at the prothallus stage grown on +1µM NAA ((e), (f)), +3µM NAA ((g), (h)) and +5µM NAA ((i), (j)) media. Elevated auxin treatment retards sporeling development with prothalli taking longer to emerge, hence this stage not being observed until 14dpg. Even with the extended growth time most sporelings grown under elevated auxin did not proceed to the prothallus stage at all. Instead, these sporelings formed large callus-type protonema (blue arrows) and did not produce any margin tissue nor exhibit any margin tissue marker signal. Those sporelings that did proceed to that prothallus stage displayed elevated levels of margin tissue marker signal, correlated with the exogenous auxin concentration. This is similar to the situation in ET239-P64 gemma grown under elevated auxin levels (see Figure S3 (a)-(g)). Scale bars= 100µm.

