## Supplementary Methods for "An Enhancer Trap system to track developmental dynamics in Marchantia polymorpha"

**Methods S1.**

**TAIL-PCR protocol for detection of enhancer trap insertion sites in the *Marchantia polymorpha* genome**

**PCR Primers**

Arbitrary Degenerate Primers

AD1	NTCGASTWTSGWGTT

AD2	NGTCGASWGANAWGAA

AD3	WGTGNAGWANCANAGA

AD4	GWGNAGSANCASAGA

AD5	AGWGNAGWANCAWAGG

AD6	STTGNTASTNCTNTGC

Specific Nested Primers

ETRB1	TGTCCAGATCGAAATCGTCTAGCGCGT

ETRB2	TCTCGAGGGAAGATCAGGAGGAACAGC

ETRB3	GGTCTTGCGAAGATCCAGTCGCTAGGT

ETLB1	TCGCAATGATGGCATTTGTAGGAGCCA

ETLB2	CGCCAGTCTTTACGGCGAGTTCTGTTA

ETLB3	GCGTAAGCCTCTCTAACCATCTGTGGG

**PCR Mix**

Primary PCR Mix

5x Phusion (Thermo Scientific) PCR Buffer 4µl

2.5mM dNTP Mix 2µl

AD Primer (Stock Concentration 20µM) 4µl

ET Primer (Stock Concentration 2µM) 2µl

Phusion (Thermo Scientific) Taq Polymerase (1U) 0.5µl

10-20ng gDNA Template

Add ddH_2_0 up to final volume of 20µl

For the primary PCR, primers ETLB1 and ETRB1 were used in all possible combinations with each of the AD primers.

Secondary PCR Mix

5x Phusion PCR Buffer 10µl

2.5mM dNTP Mix 5µl

AD Primer (Stock Concentration 20µM) 5µl

ET Primer (Stock Concentration 2µM) 5µl

Phusion (Thermo Scientific) Taq Polymerase (1U) 0.5µl

1^o^ PCR Reaction (as Template) 1µl

ddH_2_0 23.5µl

For the secondary PCR, primers ETLB2 and ETRB2 were used in all possible combinations with each of the AD primers.

Tertiary PCR Mix

5x Phusion PCR Buffer 10µl

2.5mM dNTP Mix 5µl

AD Primer (Stock Concentration 20µM) 5µl

ET Primer (Stock Concentration 2µM) 5µl

Phusion (Thermo Scientific) Taq Polymerase (1U) 0.5µl

2^o^ PCR Reaction (as Template) 1µl

ddH_2_0 23.5µl

For the tertiary PCR, primers ETLB3 and ETRB3 were used in all possible combinations with each of the AD primers.

**PCR Protocol**

Primary PCR

93^o^C 2min

(94^o^C 1min; 62^o^C 1min; 72^o^C 2min) 5 cycles

94^o^C 1min; ramp to 25^o^C (0.4^o^C/s) 3min; 25^o^C 3min; ramp to 72^o^C (0.3^o^C/s) 3min; 72^o^C 2min

(94^o^C 30s; 65^o^C 1min; 72^o^C 2min; 94^o^C 30s; 65^o^C 1min; 72^o^C 2min; 94^o^C 30s; 45^o^C 1min; 72^o^C 2min) 15 cycles

72^o^C 5min

4^o^C hold

Secondary PCR

93^o^C 1min

(94^o^C 30s; 62^o^C 1min; 72^o^C 2min; 94^o^C 30s; 62^o^C 1min; 72^o^C 2min; 94^o^C 30s; 45^o^C 1min; 72^o^C 2min) 13 cycles

72^o^C 5min

4^o^C hold

Tertiary PCR

93^o^C 1min

(94^o^C 30s; 45^o^C 1min; 72^o^C 2min) 20 cycles

72^o^C 5min

4^o^C hold
