## Supplementary material for "An Enhancer Trap system to track developmental dynamics in Marchantia polymorpha": Table S1

| Flurophore | Lines Present In | Excitation Wavelengths (nm) | Collection Wavelengths (nm) |
| --- | --- | --- | --- |
| CFP | YUCCA marker line | 442 (Diode) | 462-486 |
| eGFP | 238- lines, PIN marker line | 488 (White Light Laser) | 500-512 |
| mVenus | 238-lines, 239-lines and PIN marker line | 515 (White Light Laser) | 522-541 |
| mScarlet | 239-lines | 569 (White Light Laser) | 589-625 |
| Chlorophyll | All plants | 442 (Diode) | 670-701 |

**Table S1. Excitation and collection wavelengths used in confocal microscope imaging.**
