## Supplementary material for "An Enhancer Trap system to track developmental dynamics in Marchantia polymorpha": Legend for Movies S1, S2 and S3

**Movie S1. Time lapse video of the margin marker line ET239-P64 illustrating formation of a second row of margin tissue and z-axis split of the thallus**. Initially margin tissue signal is only present around the thallus edge. After 12 hours the first signal appears from the second, inner row of new margin tissue at the apical notch at the left of the video. By 24 hours elapsed this second row has prominent signal and by day 2 the z-axis split is clearly visible in the left apical notch. Gemma imaging began immediately after removal from the gemma cup (0dpg), and time elapsed is shown at the top. The mScarlet channel has been removed for clarity. Scale bar= 100µm.

**Movie S2. Time lapse video of the margin marker line ET239-P64 illustrating formation of a second row of margin tissue and z-axis split of the thallus**. Initially margin tissue signal is only present around the thallus edge. After 38.52 hours the first signal appears from the second, inner row of new margin tissue at the apical notch on the left of the video and appears after 43.656 hours on the apical notch on the right. By the end of the time lapse (64.2 hours elapsed) this second row has prominent signal, and the z-axis split is beginning in the apical notch on the left. Comparison with Movie S1 illustrates the variability between plants in the timing of this developmental stage, but that the formation of the second row of margin tissue precedes the z-axis split, which in turn precedes the formation of the mature, fully three-dimensional thallus with air pores and air chambers. Gemma imaging began immediately after removal from the gemma cup (0dpg), and time elapsed is shown at the top. Scale bar= 100µm.

**Movie S3. Time lapse video of the margin marker line ET239-P64 grown on media supplemented with +1µM NAA, showing the spontaneous formation of margin tissue marker signal deep in the thallus.** Abundant marker signal is present in the thallus, away from the edges and clearly inside the row of oil cells that is inside the margin tissue in normal development. New nuclear mVenus signal appears at random throughout the video in the thallus. This is not derived by production of new margin tissue from the apical notch (cf. Movie S1 and Movie S2). There is no evidence (such as new cell membrane formation or binary fission or reconstitution of fluorescent nuclei) that the new cells with marker signal are derived from existing margin tissue or thallus cells with marker signal. Instead, the gene enhancer elements trapped are being spontaneously turned on, linked to the elevated auxin treatment. Gemma was imaging began at 1dpg, and time elapsed is shown at the top. The chlorophyll channel has been removed for clarity. Scale bar= 100µm.
